# How do we imagine a speech? A triple network model for situationally simulated inner speech

**DOI:** 10.1101/2024.07.18.604038

**Authors:** Xiaowei Gao, Junjie Yang, Chaoqun Li, Xiaolin Guo, Yaling Wang, Zhiheng Qu, Wanchun Li, Jiaxuan Liu, Zhongqi Li, Wanjing Li, Zhe Hu, Junjing Li, Yien Huang, Jiali Chen, Hao Wen, Zehao Zhao, Daniel Kaiser, Tao Wei, Binke Yuan

**Author notes:** **Correspondence to:** Binke Yuan, Ph.D., Institute for Brain Research and Rehabilitation, South China Normal University, Guangzhou, China, Address: No.55, West of Zhongshan Avenue, Tianhe District, Guangzhou City. Postcode: 510631.

## Abstract

Inner speech is a silent verbal experience and plays central roles in human consciousness and cognition. Despite impressive studies over the past decades, the neural mechanisms of inner speech remain largely unknown. In this study, we adopted an ecological paradigm called situationally simulated inner speech. Unlike mere imaging speech of words, situationally simulated inner speech involves the dynamic integration of contextual background, episodic and semantic memories, and external events into a coherent structure. We conducted dynamic activation and network analyses on fMRI data, where participants were instructed to engage in inner speech prompted by cue words across 10 different contextual backgrounds. Our seed-based co-activation pattern analyses revealed dynamic involvement of the language network, sensorimotor network, and default mode network in situationally simulated inner speech. Additionally, frame-wise dynamic conditional correlation analysis uncovered four temporal-reoccurring states with distinct functional connectivity patterns among these networks. We proposed a triple network model for deliberate inner speech, including language network for a truncated form of overt speech, sensorimotor network for perceptual simulation and monitoring, and default model network for integration and ‘sense-making’ processing.

**Highlights:**

1. In ten contextual backgrounds, subjects were instructed to perform situationally simulated inner speech based on cue words.
2. The ventral parts of the bilateral somatosensory areas and middle superior temporal gyrus were as centers for seed-based co-activation pattern analyses.
3. A triple network model of language network, sensorimotor network, and default mode network was proposed for deliberate inner speech.

## 1 Introduction

People engage in imagination daily, either unconsciously or with specific intentions. Imagination encompasses various forms, including envisioning different scenarios, plans, emotional states, and speech (Hunter, 2013). The act of imagining speech, or inner speech, involves the absence of overt and audible articulation (Alderson-Day & Fernyhough, 2015; Flavell, 1999; Levine et al., 1982; Orpella et al., 2022; Perrone-Bertolotti et al., 2014). Inner speech is closely tied to thought processes and self-awareness (Morin, 2009), playing a crucial role in language monitoring, executive function, and cognitive development(Fernyhough & Borghi, 2023; Pratts et al., 2023; Alderson-Day & Fernyhough, 2015). Abnormalities in inner speech have been linked to clinical symptoms such as auditory hallucinations (Allen et al., 2007; Allen et al., 2008; Gould, 1948; Jones & Fernyhough, 2007) and rumination (Holmes et al., 2006; Holmes et al., 2009; Perrone-Bertolotti et al., 2014a).

Situationally simulated inner speech represents a distinct type of inner speech, where individuals construct scenarios and scenes within their psychological space. This form often involves dynamically imagining and articulating structured narratives, such as delivering a speech. Situationally simulated inner speech involved (1) a process of inner speech, (2) the integration of contextual background, episodic and semantic memories, past experiences, and upcoming events to construct contextually coherent narratives (Andrews, 2016, 2014; Kaiser et al., 2022), and (3) the representation of complex and dynamic information, contrasting with simple imagining of vowels, consonants, tones, and words. Despite extensive research on the neural mechanisms of inner speech more generally (Alderson-Day & Fernyhough, 2015; Fernyhough & Borghi, 2023; Grandchamp et al., 2019; Perrone-Bertolotti et al., 2014 ), the neural mechanisms underlying situationally simulated inner speech remain largely unknown.

As a distinct form of inner speech, situationally simulated inner speech likely shares some neural mechanisms with other forms of inner speech. Extensive research indicates that inner speech activates specific brain regions, including the left inferior frontal gyrus (Broca’s area) (Aleman et al., 2005; Hurlburt et al., 2016; Shergill et al., 2001), the supplementary motor area, the premotor cortex, and the insular cortex (Aleman et al., 2005; Cooney et al., 2018; Shergill et al., 2001). These regions are primarily associated with motor planning during typical speech production (Bohland & Guenther, 2006; Guenther, 2006; Guenther et al., 2006; Guenther & Vladusich, 2012; Kearney & Guenther, 2019). Additionally, inner speech has been found to engage the cerebellum (Ackermann & Hertrich, 2003; Naito et al., 2002), which plays a crucial role in motor planning for articulation (Kearney & Guenther, 2019; Parrell et al., 2017). Inner speech, encompassing both “articulation imagery” rooted in somatosensory-motor processes and “hearing imagery” derived from memory recall, engages the temporal lobe cortex, as evidenced by several studies (Courson & Tremblay, 2020; Sereshkeh et al., 2018; Tian & Poeppel, 2012; Tian et al., 2016). This suggests that inner speech involves not only simulating speech production but also sensorimotor processes, integrating articulatory and sensory representations. Hence, we propose that situationally simulated inner speech necessitates a dynamic recruitment of the sensorimotor network.

Moreover, as a multifaceted and dynamic phenomenon, situationally simulated inner speech demands the amalgamation of information across various time scales, including phonemes, morphemes, words, sentences, actions, roles, events, and narrative frameworks (Sonkusare et al., 2019; Willems et al., 2020). Numerous functional neuroimaging investigations have highlighted the involvement of the default mode network (DMN) in this “meaning-making” function (Yeshurun Y, 2021), particularly in the integrative processing of events (Buckner & DiNicola, 2019; Chang et al., 2021; Roger et al., 2022). Notably, DMN exhibits positive activations during narrative comprehension (Menon, 2023; Simony et al., 2016; Song et al., 2021; Yeshurun et al., 2021). The computational processes underlying integration encompass extraction/dimensionality reduction, multimodal/relationship binding, and pattern separation/completion (Cowell et al., 2019; Roger et al., 2022). Therefore, we also hypothesized that situationally simulated inner speech requires the involvement of DMN.

To validate these hypotheses, we conducted dynamic activation and connectivity analyses on an fMRI dataset (Kaiser et al., 2022) comprising 10 sessions of situationally simulated inner speech. In each session, participants were presented with 61 abstract words and instructed to mentally incorporate them into a cohesive narrative within a predefined contextual backdrop. For example, participants imagined themselves as astronauts exploring distant galaxies, endeavoring to elucidate humanity and its culture to an enigmatic alien species. This task necessitated not only retrieving word meanings but also employing them to weave into a complex ongoing stream of thought. To unveil the transient brain activations and network reconfigurations during these processes, we employed co-activation pattern (CAP) analysis (Bolton et al., 2020; Liu & Duyn, 2013; Liu et al., 2018) and dynamic conditional correlation (Aielli, 2013; Engle, 2002; Yuan et al., 2023a, 2023b). The core brain regions utilized for CAP analysis were derived from a term-based meta-analyses of 14 inner-speech-related terms, and further corroborated through.an independent fMRI dataset involving covert speech of words and numbers.

## 2 Materials and Methods

### 2.1 Term-based meta-analysis

To elucidate the core brain regions engaged in inner speech, we conducted term-based meta-analyses. Initially, we compiled key terms associated with inner speech from Alderson-Day and Fernyhough’s work, encompassing keywords such as inner speech, private speech, self-talk, covert speech, silent speech, verbal thinking, verbal mediation, inner monologue, inner dialogue, inner voice, articulatory imagery, voice imagery, speech imagery, and auditory verbal imagery (Alderson-Day & Fernyhough, 2015). Subsequently, for each term, we performed a meta-analysis using Neuroquery (https://neuroquery.org/) (Dockès et al., 2020), an automated meta-analysis tool for neuroimaging research. Neuroquery incorporates 13,459 neuroimaging publications and 7,547 neuroscience terms. Unlike conventional statistical-based meta-analysis methods, NeuroQuery employs supervised machine learning to predict a brain map, particularly useful for terms with limited study or arbitrary text lengths. Leveraging this meta-analytic exploratory tool, we obtained 14 activation maps. To identify consistent activations across terms, we initially thresholded the 14 activation maps (Z = 3, cluster size = 20) and then aggregated them.

### 2.2 Localization of core brain regions in inner speech

To further validate the results of the core brain regions of inner speech, we sought to independently derive a set of regions of interest that are related to inner speech. The fMRI dataset utilized for identifying the core brain regions involved in inner speech was reanalyzed from the “Bimodal” dataset (Simistira Liwicki et al., 2023). This dataset includes publicly available EEG and fMRI data acquired during covert speech of words and numbers. Five healthy right-handed participants (mean age 39 years, SD = 7.07; three females) participated in this study.

The fMRI experiment comprised two sessions conducted over three days. Word stimuli for the experiment included social category words (such as child, daughter, father, and wife) and number category words (such as four, three, ten, and six), with each category containing four words, totaling eight different words. Word stimuli were presented in a random order. Each session commenced with a 2-second fixation period. Subsequently, each trial consisted of an inner speech task lasting 4 seconds, followed by a rest period of 10 seconds. During the inner speech task, word stimuli were continuously presented for 4 seconds, and participants were instructed to mentally repeat the given word as many times as possible (approximately 4 times) without any accompanying pronunciation or muscle movement. Throughout the rest period, a white fixation cross was displayed for 10 seconds, allowing participants to relax and prepare for the subsequent trial. The inner speech task incorporated eight different words, categorized into social or number words, with each word comprising 20 trials per session. Consequently, each session consisted of 160 trials, and the total recording time for 320 repetitions per participant was 74.6 minutes. This is depicted in Supplementary Figure 1. Due to significant fluctuations in sub-04’s during the EEG recording, the “Bimodal” dataset did not include data from this participant.

Functional MRI data was acquired using the Siemens Magnetom Prisma MRI system equipped with a 20-channel head coil. Anatomical images were obtained using a sagittal T1-weighted 3D magnetization-prepared rapid gradient-echo (MPRAGE) sequence with the following parameters: repetition time (TR) = 2300 milliseconds; echo time (TE) = 2.98 milliseconds; inversion time (TI) = 900 milliseconds; flip angle = 9 degrees; number of slices = 208; matrix size = 256×256; voxel size = 1×1×1 millimeters. Immediately after the anatomical scan, two field maps (A and B) were acquired with the following parameters: TR = 662.0 milliseconds; echo time (TE): A = 4.92 milliseconds, B = 7.38 milliseconds; voxel size = 3 × 3 × 2 millimeters. Subsequently, functional magnetic resonance imaging was performed using an echo-planar imaging sequence with multiband acceleration factor = 2 and in-plane acceleration factor = 2, parallel to the double oblique plane, with the following parameters: TR = 2.16 seconds; TE = 30 milliseconds; number of slices = 68; matrix size = 100×100; voxel size = 2×2×2 millimeters.

We preprocess the publicly available raw data downloaded using SPM12 (www.fil.ion.ucl.ac.uk/spm/), following the processing steps and parameter settings as described in the original article (Simistira Liwicki et al., 2023).

To identify the core brain regions implicated in inner speech, individual-level analyses were performed utilizing the General Linear Model (GLM) in SPM12 software, based on the Matlab R2020a platform (http://www.fil.ion.ucl.ac.uk/spm). Category regressors (social or number words) were synchronized with the onset of their respective inner speech words and persisted for 4 seconds. The remaining regressors lasted for 10 seconds. Additionally, six head motion parameters computed during motion correction were included as covariates. For each condition, regressors were modeled as boxcar functions and convolved with the standard hemodynamic response function (Friston et al., 1994). A high-pass filter was set to 128 seconds. Voxel-wise thresholding was executed with False Discovery Rate (FDR) correction, employing a voxel-level threshold of ^K^E > 20, corrected p < 0.05. Due to the limited number of participants, group-level analyses were not conducted.

### 2.3 fMRI dataset of a situationally simulated inner speech paradigm

After identifying the core regions involved in inner speech, we investigated the activation dynamics among these regions in a dataset that involved situationally simulated inner speech (Kaiser et al., 2022). This dataset comprises ten fMRI runs involving 19 healthy participants (mean age 28.8 years, SD = 6.1; 10 females). Two participants were excluded as they completed a different version of the experiment with a 1s inter-trial interval instead of 1.5s and did not include the behavioral task.

The stimulus set, consisting of 61 German abstract nouns selected from the most frequently used German word list (selected from: wortschatz.uni-leipzig.de), covered various topics (Supplementary Table 1). During the fMRI experiment, participants completed 10 runs. Prior to each run, participants read one of the 10 background stories (Supplementary Table 2) and were instructed to mentally immerse themselves in the outlined situation. Subsequently, participants silently crafted a narrative in their minds using the sequentially presented words on the screen (Figure 1). They were informed that the story’s progression relied solely on their utilization of all the words. Selected stories were emotionally engaging to enhance participants’ task immersion. The order of the 10 stories was randomized for each participant, with each run containing 61 experimental trials. In each trial, each word was displayed for 3 seconds with a 1.5-second interval, during which a fixation cross was presented. Additionally, each run incorporated 14 randomly presented fixation trials displayed throughout the experiment. The trial order was randomized within each run. To maintain participants’ attention, seven words were presented in pink, prompting participants to press a button upon sighting them. Brief fixation periods were included at the start and end of each run, with each run lasting 5 minutes and 48 seconds.

**Figure 1.**
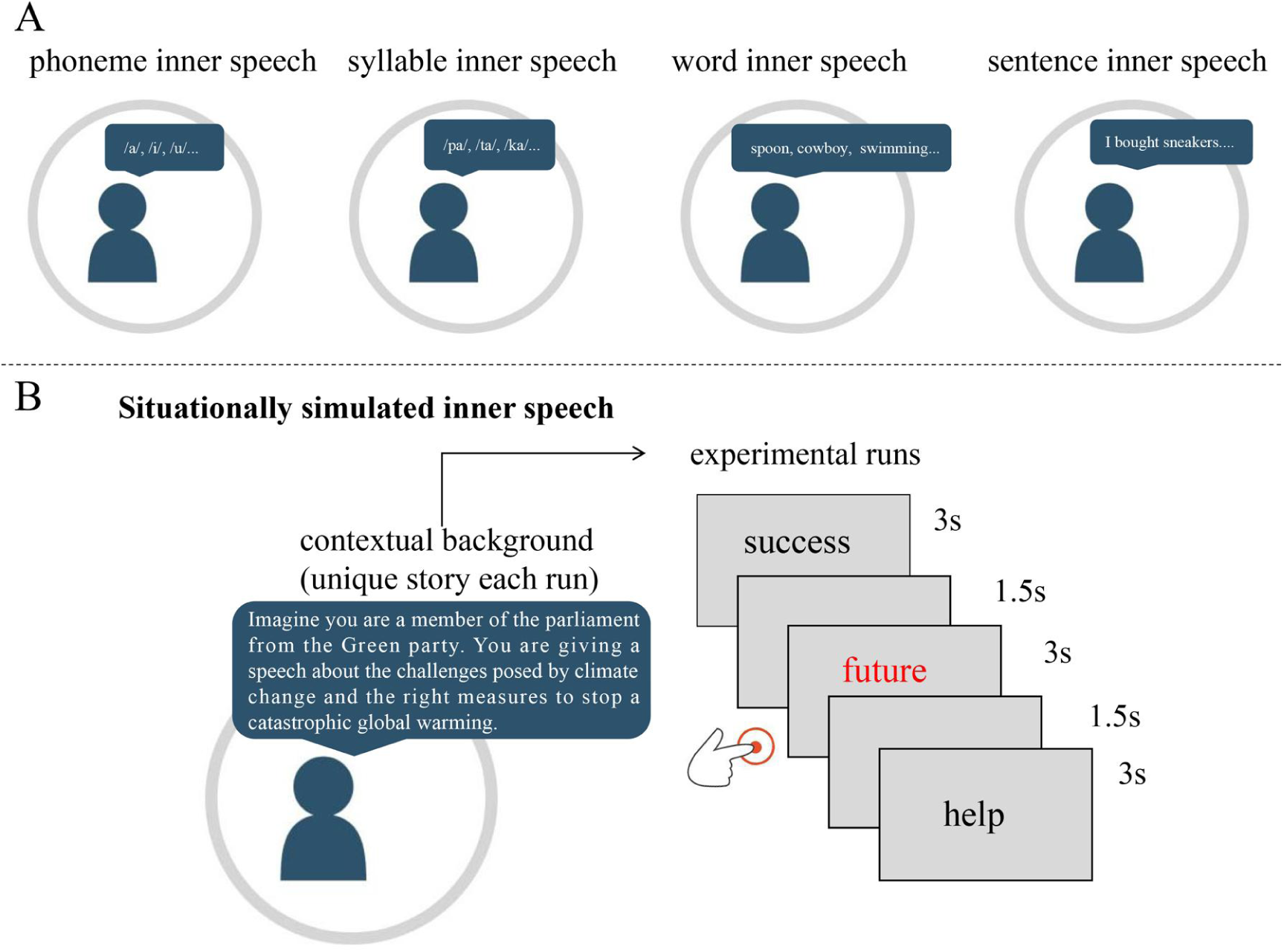
A: Typical methods for eliciting inner speech involve covertly pronouncing different phonemes, syllables, words, and even sentences. B: Experimental design for situationally simulated inner speech. Each participant completed 10 fMRI recording runs. Before each run, participants mentally constructed a unique background context and were instructed to silently narrate a story incorporating 61 abstract words. During each trial, individual words were displayed for 3 seconds, separated by a 1.5-second interval, with a fixation cross shown in between. To maintain participants’ attention, 7 words were presented in pink, prompting participants to press a button. Cue words were presented in a random order during each run. English translations for the 61 abstract words can be found in Supplementary Table 1.

MRI data was acquired using a 3T Siemens Tim Trio Scanner equipped with a 12-channel head coil. T2-weighted gradient-echo echo-planar images were collected as functional volumes (TR = 2s, TE = 30ms, 70 °flip angle, 3mm × 3mm voxel size, 37 slices, 20% gap, 192mm FOV, 64×64 matrix size, interleaved acquisition). Additionally, a T1-weighted image (MPRAGE; 1mm × 1mm × 1mm voxel size) was obtained as a high-resolution anatomical reference.

Data preprocessing was performed in MATLAB using SPM12 (www.fil.ion.ucl.ac.uk/spm/) and DPABI (http://rfmri.org/dpabi). The preprocessing steps included: 1) slice timing correction; 2) head motion correction. No participant had head motion exceeding 3 mm maximum translation or 3°maximum rotation; 3) coregistration of the motion-corrected functional images to the 3D-T1 image, followed by spatial normalization to MNI space using the estimated deformation field from segmentation; 4) spatial smoothing of the normalized images using a Gaussian kernel (full width at half maximum = 6 mm); 5) detrending; 6) regression of nuisance signals. Twenty-four parameters were removed from each voxel’s time series using linear regression, including six parameters of translation and rotation, their time derivatives (6 parameters), and quadratic terms (12 parameters) (Yan et al., 2013); 7) temporal band-pass filtering (0.01-0.1 Hz).

#### 2.3.1 Brain activations using GLM

Brain activations associated with external stimuli (i.e., word processing) and subject responses (i.e., finger pressing) not relevant to this study were identified using the General Linear Model (GLM) in SPM12. Individual-subject analyses were performed by constructing a general linear model for each condition. Two regressors were defined: word and response. The six head motion parameters computed during realignment were included as covariates. For each condition, the regressor was modeled as box-car functions and convolved with a canonical hemodynamic response function (Friston et al., 1994). The high-pass filter was set at 128 s. For the second level, one-sample T-tests were used to assess main effects of experimental conditions. In all analyses, significant voxel clusters on each t-map were identified with FDR correction at p < 0.05.

#### 2.3.2 Seed-based co-activation pattern analysis

To delineate the networks engaged in situationally simulated inner speech, we conducted seed-based co-activation pattern analysis (CAP). The core brain regions of inner speech identified through analysis of the “Bimodal” dataset and term-based meta-analysis were utilized as seeds.

CAP is a frame-wise method for assessing instantaneous whole-brain activation patterns (Bolton et al., 2020; Liu & Duyn, 2013; Liu et al., 2018). It operates on the premise that high-amplitude fluctuations (co-activation or de-activation) drive functional connectivity and networks (Tagliazucchi et al., 2012; Zamani Esfahlani et al., 2020). Essentially, it involves extracting frames at time points with suprathreshold signals, followed by averaging or clustering these signals to represent temporal recurring patterns. In seed-based correlation analysis, the network patterns associated with a given seed are typically estimated by linearly correlating the time series of each gray matter voxel with the referenced seed. The CAP method (Liu & Duyn, 2013) demonstrates that identical network patterns can be obtained by voxel-wise averaging of spatial maps from time frames where the seed signal intensity surpasses a certain threshold. Temporal clustering of these extracted time frames based on their spatial similarity can generate multiple spatial patterns, conjectured to be functionally relevant and reflective of co-activation patterns (CAPs) across the whole brain at individual time frames (Chen et al., 2015).

We employed the TbCAPs toolbox (https://c4science.ch/source/CAP_Toolbox.git) (Bolton et al., 2020) for seed-based CAP analysis. Initially, based on the results of activation analysis and meta-analysis, bilateral sensorimotor ventral parts and bilateral superior temporal gyri were selected as seed points. To mitigate motion effects, scrubbing was performed to retain only frames where framewise displacement did not exceed a threshold of 0.5 mm. CAPs were generated by applying k-means clustering to data from the entire gray matter volume based on thresholded frames from the seeds. Since k-means clustering is an iterative process with no guaranteed convergence towards the global optimum, the algorithm was run 50 times. The optimal number of clusters (K) was determined using consensus clustering from 2 to a user-specified Kmax. The optimal K was determined as the one minimizing the proportion of ambiguously clustered pairs (PAC), i.e., pairs that are sometimes clustered together and sometimes separately. Stability was defined as 1 - PAC. Once the optimal K was determined, k-means clustering was performed using that cluster number on all data. The percentages of positive- and negative-valued voxels to retain in each frame for clustering were set at 100%, and 100%, respectively. The threshold value for selecting (de)active frames was set at 1. The distance measure used for clustering was spatial correlation.

To assess the spatial similarities of CAPs across the 10 runs, Pearson correlation coefficients were calculated between each pair of identified thresholded CAPs maps (Z > 1, cluster size = 20 voxels).

### 2.4 Dynamic Conditional Correlation (DCC)

To explore the dynamic functional interactions among co-activation brain regions implicated in situationally simulated inner speech, we employed dynamic conditional correlation (DCC) analysis (Lindquist et al., 2014) to construct the framewise functional connectivity network. DCC has demonstrated superior performance over the standard sliding-window approach in tracking network dynamics (Choe et al., 2017; Lindquist et al., 2014). We conducted DCC analyses by using the DCC toolbox (https://github.com/caWApab/Lindquist_Dynamic_Correlation/tree/master/DCC_tool box).

The core idea of DCC is to represent the conditional variances of individual time series at a specific moment as a linear combination of past conditional variances and past time series values. It involves two steps. Firstly, standardized residuals of each time series are estimated using a univariate GARCH (1,1) process. Secondly, an exponentially weighted moving average (EWMA) window is applied to the standardized residuals to compute a non-normalized version of the time-varying correlation matrix between Regions of Interest (ROIs). Mathematical expressions of the GARCH (1,1) model, DCC model, EWMA, and estimations of the model parameters can be found in (Lindquist et al., 2014) and related literature (Yuan, Xie, & Gong et al., 2023; Yuan, Xie, & Wang et al., 2023b).

For each run, frame-wise DCC yielded 175 (175 volumes) functional connectivity matrices. To identify temporally reoccurring states, the k-means clustering algorithm was employed to decompose all participants’ networks into several non-overlapping clusters. The number of clusters (k) was determined using the elbow criterion of the cluster validity index, computed as the ratio between within-cluster distance to between-cluster distance. Subsequently, spatiotemporal characteristics were calculated, including nodal degree, overall state frequency, and state transition vectors (Yuan et al., 2023). The nodal degree for a given node was calculated by summing up the correlation coefficients of all suprathreshold connections connected with this node. Before calculation, every matrix was thresholded (r > 0.2) to eliminate weak and non-significant correlations possibly arising from noise. Given the debatable nature of negative connections (Murphy & Fox, 2017), only positive connections were considered in this work. Overall state frequency refers to the percentage of a state occurring in all frames. For each run, we concatenated all subjects’ window membership vectors and computed the percentage of the total numbers of a given state to the total number of windows. Itis important to note that the overall state frequency is proportional to the total dwell times of a state in all subjects.

To assess the spatial similarities of time-recurring states among the 10 runs, we calculated Pearson correlation coefficients between each pair of states.

## 3 Results

### 3.1 Results of term-based meta-analyses

As depicted in Figure 2, term-based meta-analyses showed strong activation in the bilateral superior temporal gyri (STG) and ventral sensorimotor areas, suggesting their potential role as core brain regions for inner speech. Specifically, the meta-analysis results demonstrated activation in the bilateral STG in 11 out of 14 terms, and activation in the ventral sensorimotor area in eight terms. Together with our analysis on the “Bimodal” dataset (Simistira Liwicki et al., 2023), these results provide a critical benchmark for the brain network involved in inner speech, which we applied during the analysis of situationally simulated inner speech in the following.

**Figure 2.**
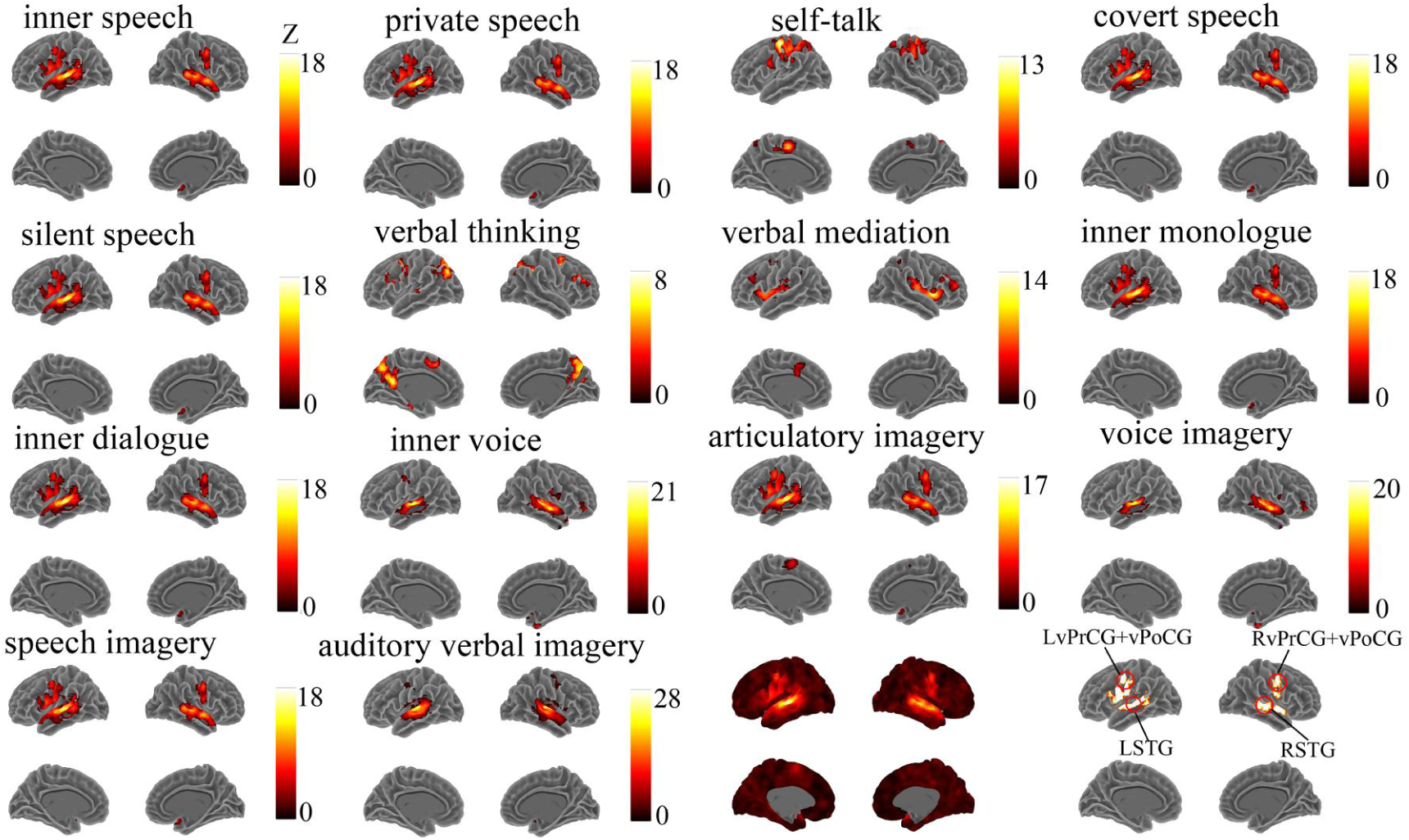
Term-based meta-analysis results of inner speech. Term-based meta-analysis results are displayed after applying a 20-voxel threshold and a threshold of 3 (a typical value used in NeuroSynth). The raw and thresholded (bilateral STG Z > 70 and bilateral ventral sensorimotor area Z > 50) overlap maps associated with inner speech are shown. Strong activations were observed in the bilateral superior temporal gyri and ventral sensorimotor areas.

### 3.2 Brain activations of speaking words and numbers covertly

To identify the core brain regions involved in inner speech, we first analyzed data from an unrelated fMRI dataset investigating the neural correlates of inner speech (Simistira Liwicki et al., 2023). As depicted in Supplementary figure 2, compared to the baseline resting condition, the inner speech task elicited activation in bilateral ventral precentral gyri, Wernicke’s area in the left hemisphere, and Broca’s area and surrounding regions. Additionally, activation was observed in the left supramarginal gyri and angular gyri, associated with semantic processing (Devlin et al., 2003; Hartwigsen et al., 2016). Notably, visual processing areas also showed heightened activation, possibly linked to the processing of visual word forms (Simistira Liwicki et al., 2023).

### 3.3 Brain activations of word processing and button pressing in situationally simulated inner speech

Next, we turned to the fMRI dataset on situationally simulated inner speech (Kaiser et al., 2022). In a first analysis, we assessed which regions show above-baseline activation related to word reading and task-related button presses. Consistent with previous studies, visual word processing activated the visual word form area (VWFA), left superior temporal gyri, and inferior frontal gyri (Supplementary figures 3A and C). The right-hand finger button press activated the left posterior central gyri and right angular gyri. The activation pattern for visual word processing and button responses remained consistent across the 10 background runs (Supplementary figures 3B and D).

### 3.4 Seed-based Co-activation Pattern Analysis (CAP) Results

For the CAP analysis, we derived seed regions from the inner speech network previously identified in the “Bimodal” dataset and the term-based meta-analysis. Adopting the ventral part of the bilateral precentral gyri and postcentral gyri (vPreCG+vPoCG) as seed points in CAP, the optimal number of clusters (K) was 3. The co-activation patterns of the left and right vPreCG+vPoCG were highly consistent. The first CAP co-activated with vPreCG+vPoCG included the sensorimotor network (SMN), encompassing the primary motor cortex, precentral gyri, postcentral gyri, supplementary motor area, and primary somatosensory cortex, as well as the visual network (VN). The second CAP primarily involved the seed points themselves. The third CAP mainly included the seed points and the language network (LAN) (Figures 3 and 4). LAN predominantly consists of the left superior temporal gyri, superior temporal gyri, inferior frontal gyri, and ventral precentral gyri, as documented by various studies (Lipkin et al., 2022; Malik-Moraleda et al., 2022; Yuan, Xie, & Wang et al., 2023c). A high spatial similarity was observed between the time-recurring states across the 10 story scans (Supplementary Figure 4). Temporally, these three states appeared at different time points and exhibited some inconsistency between participants (Supplementary Figures 5 and 6).

**Figure 3.**
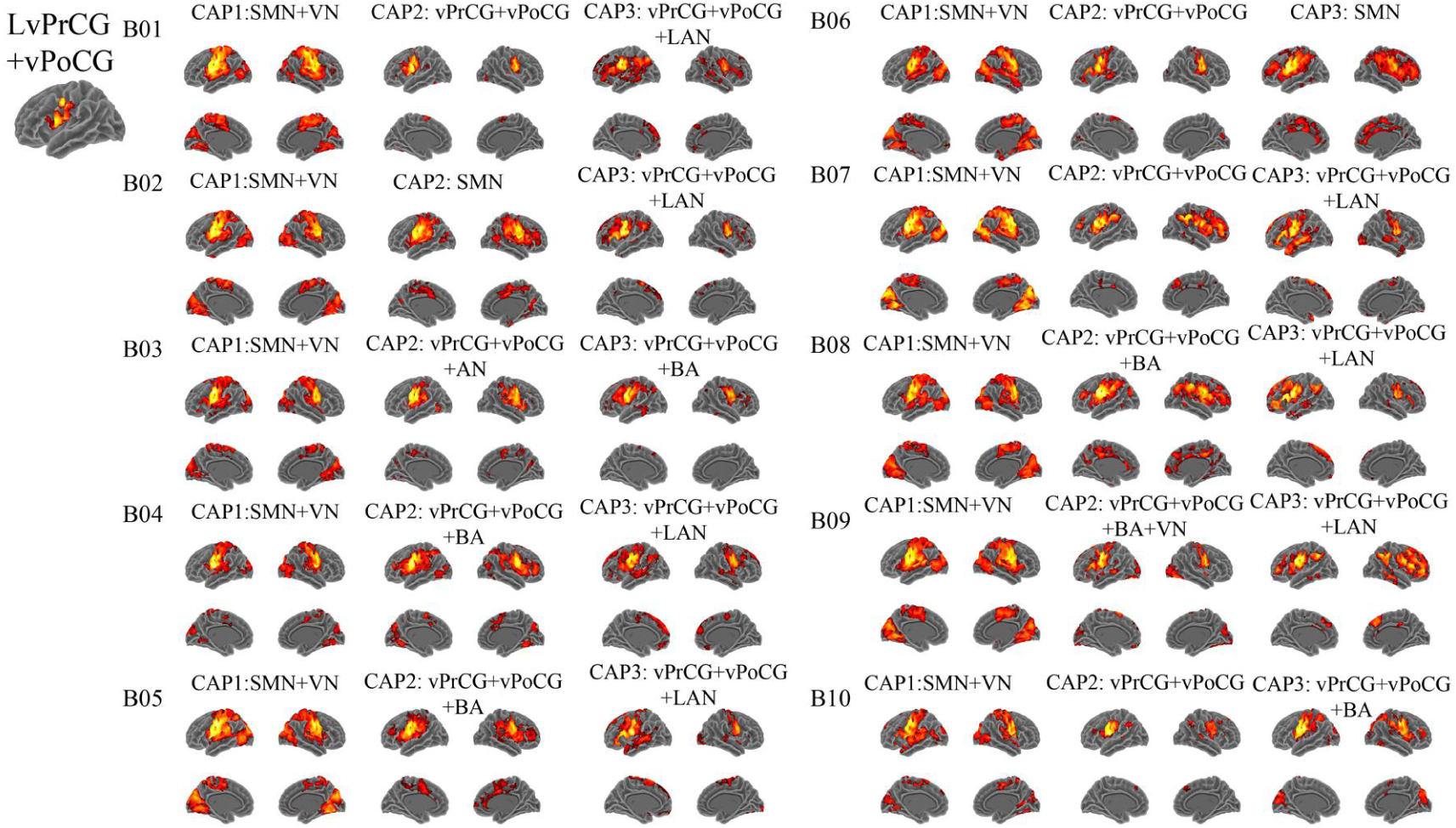
Co-activation patterns during the situationally simulated inner speech task with the ventral part of the left precentral gyri (PrCG) and postcentral gyri (PoCG) (Z > 1, cluster size = 20 voxels) across the 10 background stories. (SMN: sensorimotor network; AN: auditory network; VN: visual network; LAN: language network; BA: Broca’s area; B: background)

**Figure 4.**
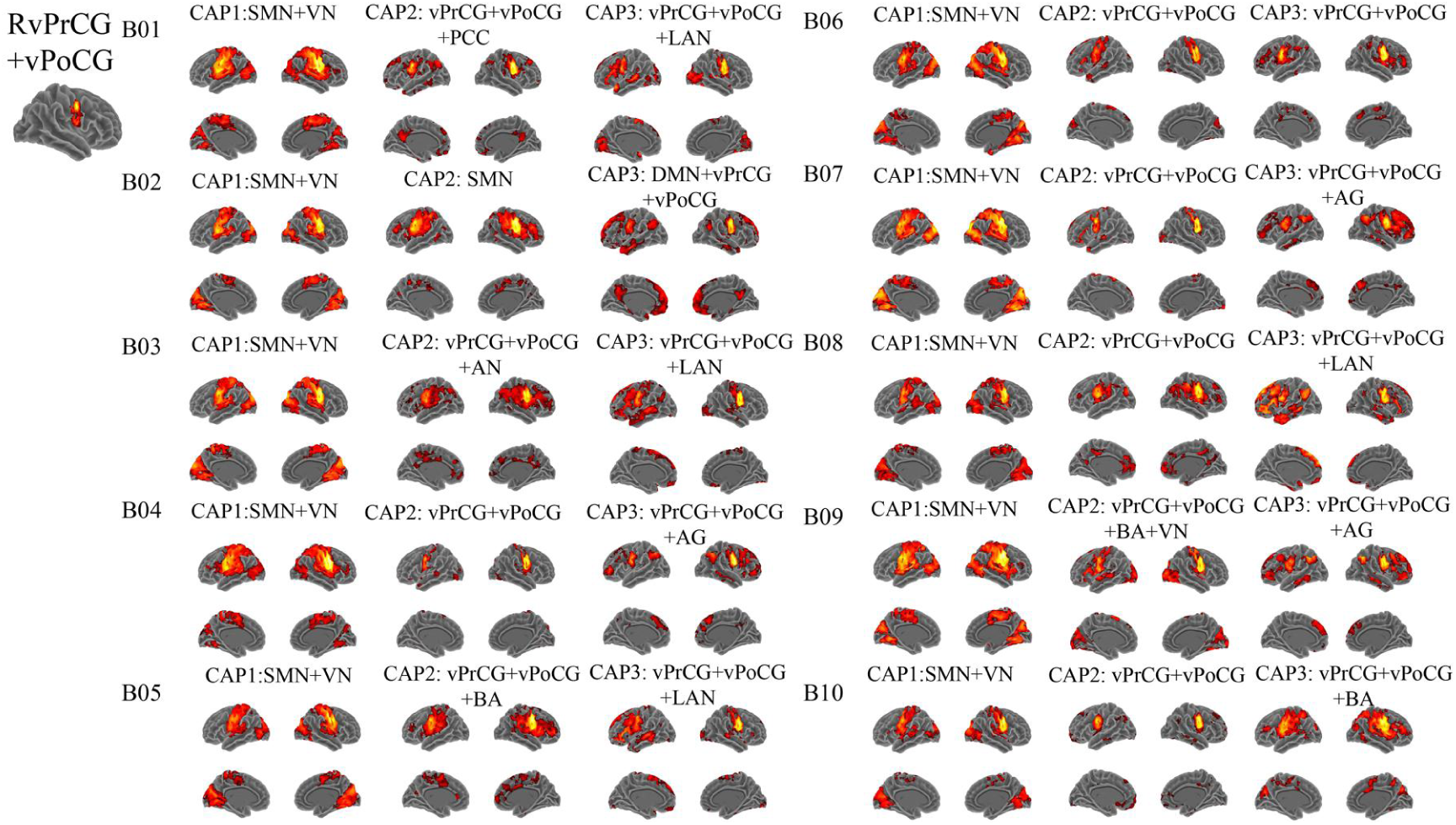
Co-activation patterns during the situationally simulated inner speech task with the ventral part of the right precentral gyri and postcentral gyri (Z > 1, cluster size = 20 voxels) across the 10 background stories. (SMN: sensorimotor network; AN: auditory network; VN: visual network; LAN: language network; BA: Broca’s area; PCC: posterior cingulate cortex; B:background)

In addition, three co-activation patterns (CAPs) were observed to co-activate alongside the bilateral superior temporal gyrus (STG). Notably, the co-activation patterns of the left and right STG exhibited remarkable consistency. These CAPs manifested as follows: the sensorimotor network, the language network, and the default mode network (posterior cingulate cortex, precuneus, and angular gyri) (Figures 5 and 6). Furthermore, a high spatial similarity is observed among time-recurring states across the ten story scans (Supplementary Fig 7). Temporally, the emergence of the three CAPs varies across different time points and again shows inconsistency between participants (Supplementary figures 8 and 9).

**Figure 5.**
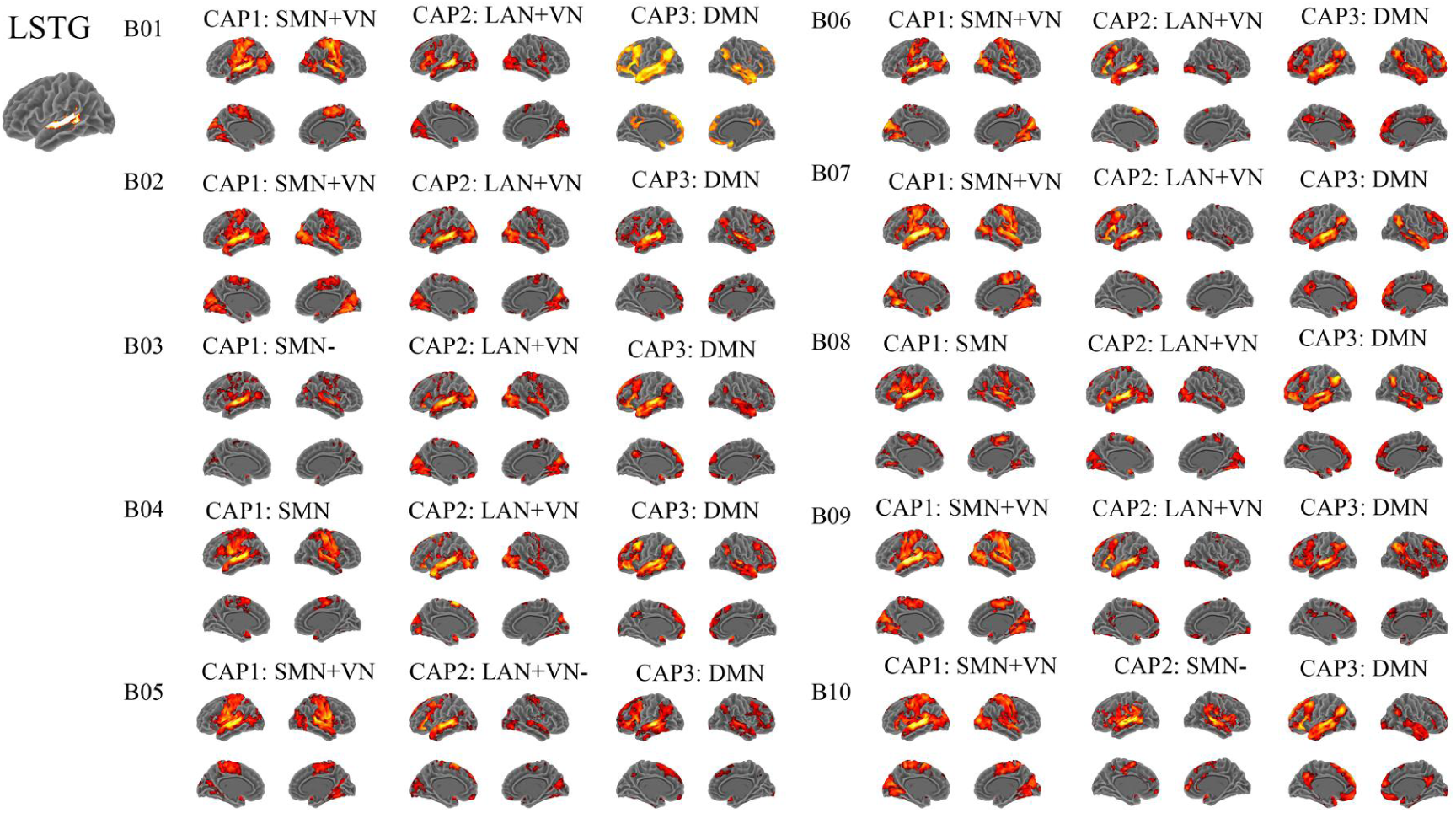
Co-activation patterns during the situationally simulated inner speech task with the left superior temporal gyri (Z > 1, cluster size = 20 voxels) across the 10 background stories. (SMN: sensorimotor network; VN: visual network; LAN: language network; DMN: default mode network; B:background)

**Figure 6.**
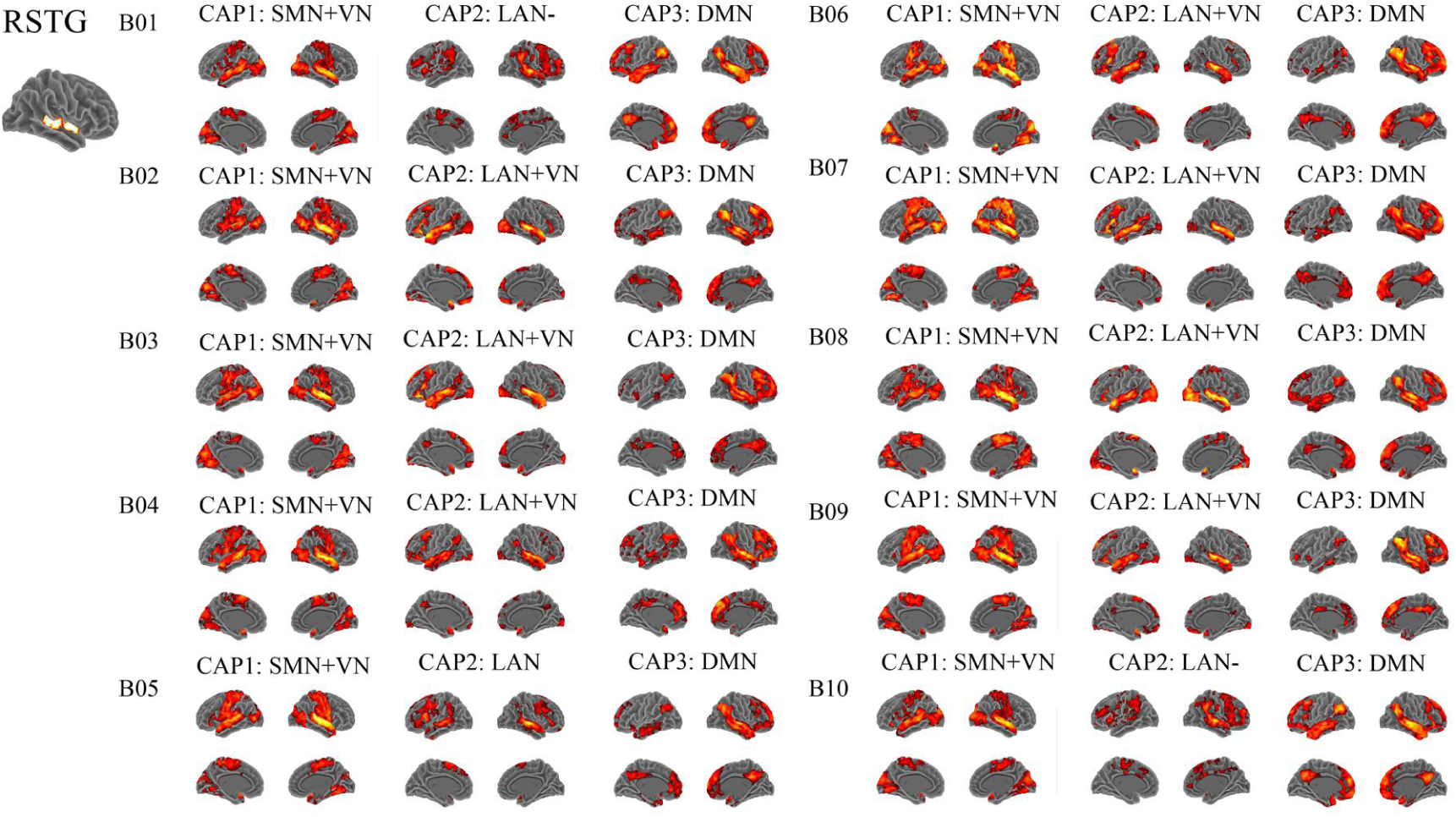
Co-activation patterns during the situationally simulated inner speech task with the right superior temporal gyri (Z > 1, cluster size = 20 voxels) across the 10 background stories. (SMN: sensorimotor network; VN: visual network; LAN: language network; DMN: default mode network; B:background)

In the CAP results, we also noted substantial activations in cerebellar regions (Supplementary Figures10-11), specifically in regions 7, 8, and 9 as delineated by the multi-domain task battery parcellation (Gaiser et al., 2024).

### 3.5 Frame-wise dynamic functional network reorganization

Based on the co-activation patterns of the ventral parts of the bilateral precentral gyri and postcentral gyri, we conducted Dynamic Conditional Correlation (DCC) analysis using nodes defined by the Brainnetome Atlas (Fan et al., 2016), representing the language network (LAN) and sensorimotor network (SMN). This analysis produced a functional connectivity matrix consisting of 69 nodes (41 nodes related to LAN and 28 nodes related to SMN). By applying the k-means clustering algorithm and the elbow criterion to the matrix of each story (69 x 69 x 175 x 17), we identified four distinct time-recurring states (Figures 7-8). Robust functional connections were observed within LAN and SMN, as well as between LAN and SMN. In State 2, strong connections were noted within SMN and among some nodes of LAN. State 3 revealed strong connections within LAN, particularly among nodes in the inferior frontal gyri (IFG). In State 4, overall connectivity patterns appeared weaker. Across all states, consistent strong connections within the IFG and STG were observed, indicating sustained engagement of situationally simulated inner speech by participants. Notably, the time-recurring states exhibited high similarity across the 10 background stories and demonstrated high spatial similarity (Supplementary figure 12). State 4 exhibited the highest overall frequency across the 10 background stories (Supplementary figure 13).

**Figure 7.**
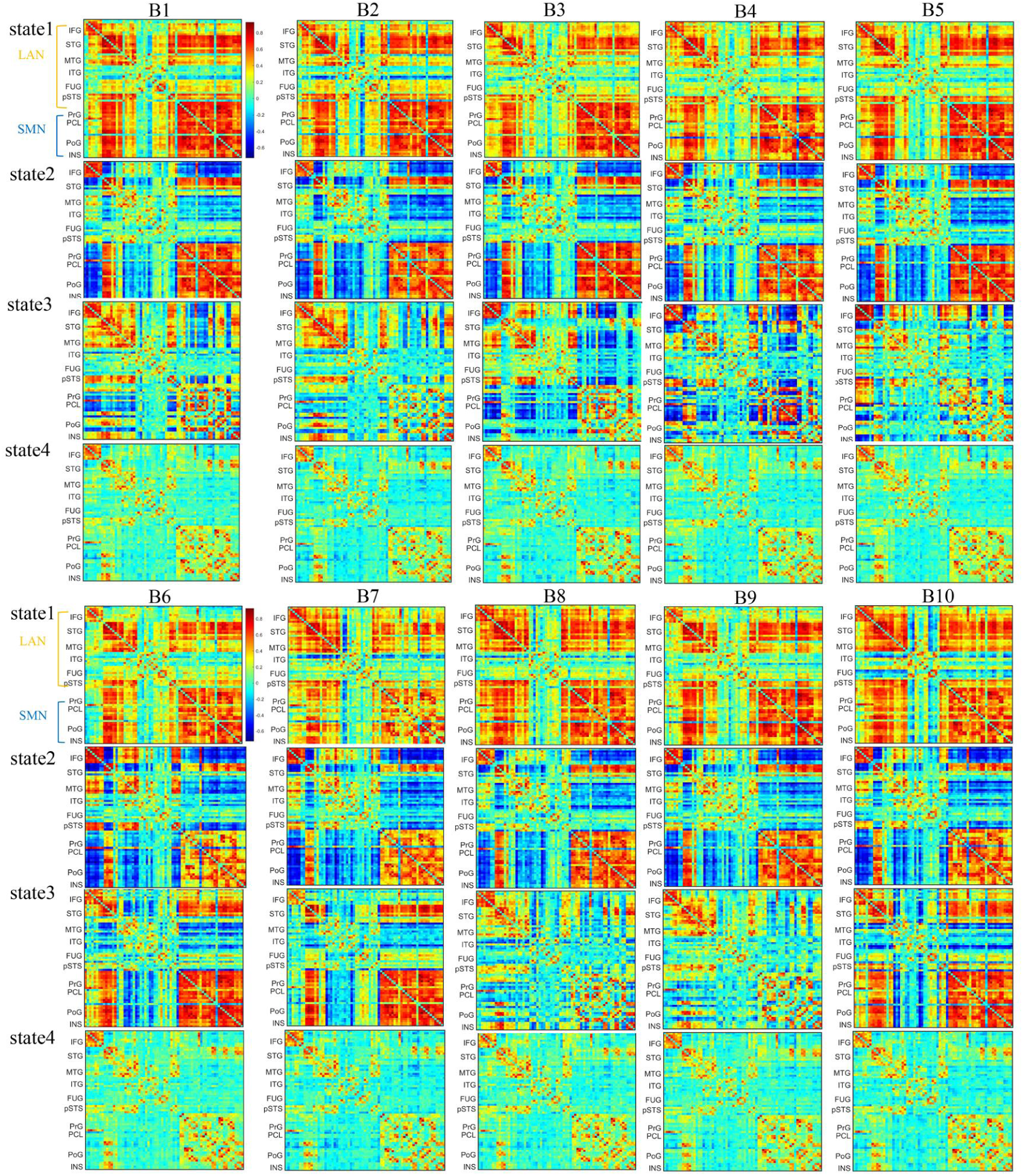
Four states identified by DCC through K-means clustering in two functional networks across background stories 1 to 10. Each functional connectivity matrix illustrates the intensity of functional connections between various nodes. (B:background)

**Figure 8.**
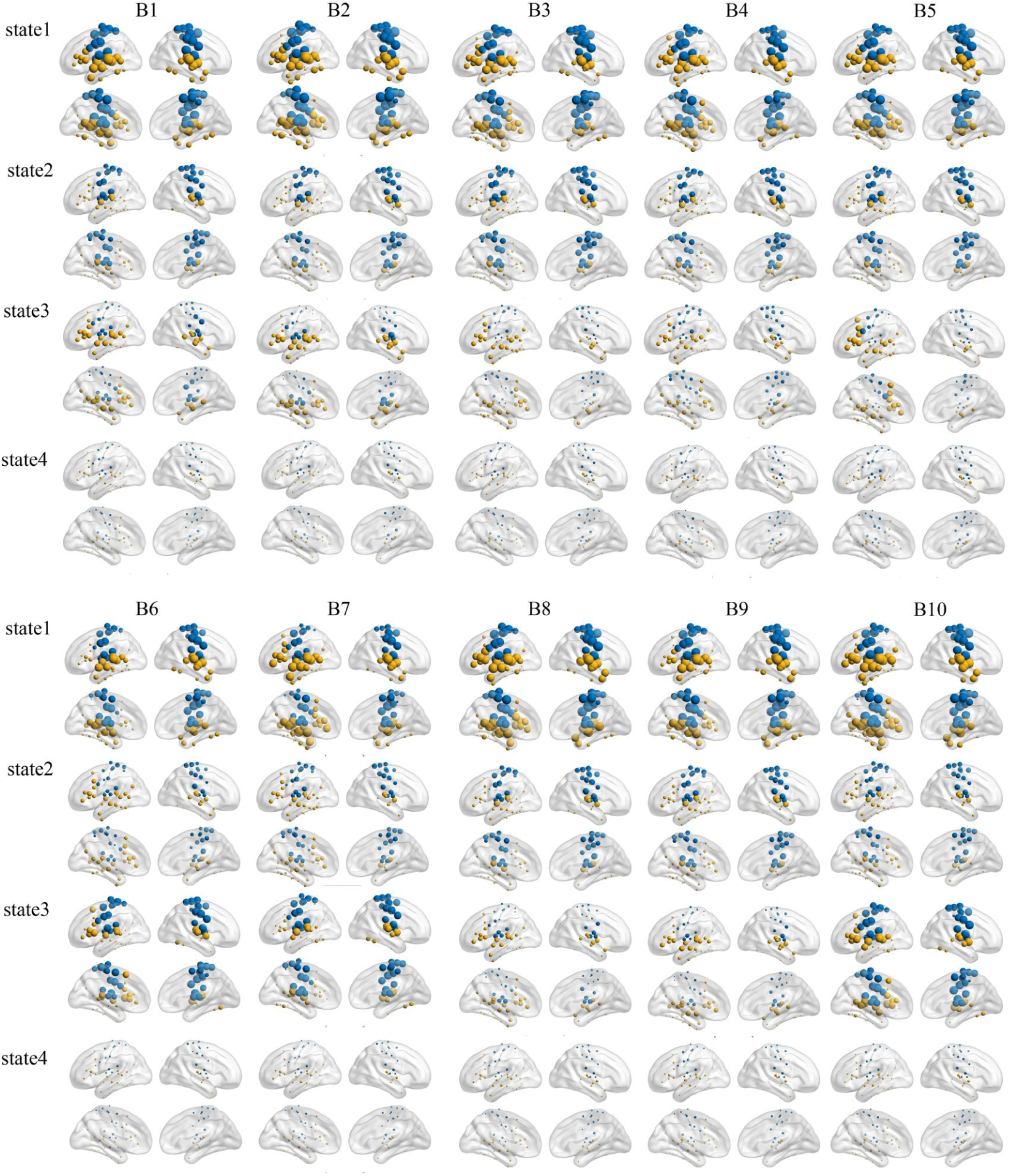
Activation node degree of DCC in two functional networks across background stories 1 to 10. The color of each node indicates its affiliation with different brain networks (blue: sensorimotor network; yellow: language network; red: default mode network), while the size of the node represents its activation intensity. (B:background)

Building upon the co-activation patterns of the bilateral middle temporal gyri, we extended our investigation to include the default mode network (DMN) alongside LAN and SMN. This analysis yielded a functional connectivity matrix comprising 94 nodes (41 LAN nodes, 25 DMN nodes, and 28 SMN nodes). Through the application of the k-means clustering algorithm and the elbow criterion to the matrix of each story (94 x 94 x 175 x 17), we discerned four distinct time-recurring states (Figure 9-10). The DCC activation results paralleled those derived from the two functional networks mentioned above. However, in State 2, a significant antagonistic interaction emerged between SMN and DMN. State 3 showcased robust internal correlations within the DMN. Notably, the time-recurring states exhibited high similarity across the 10 background stories and demonstrated a high degree of spatial congruence (Supplementary figure 14). Once again, State 4 emerged with the highest overall frequency across the 10 background stories (Supplementary figure 15).

**Figure 9.**
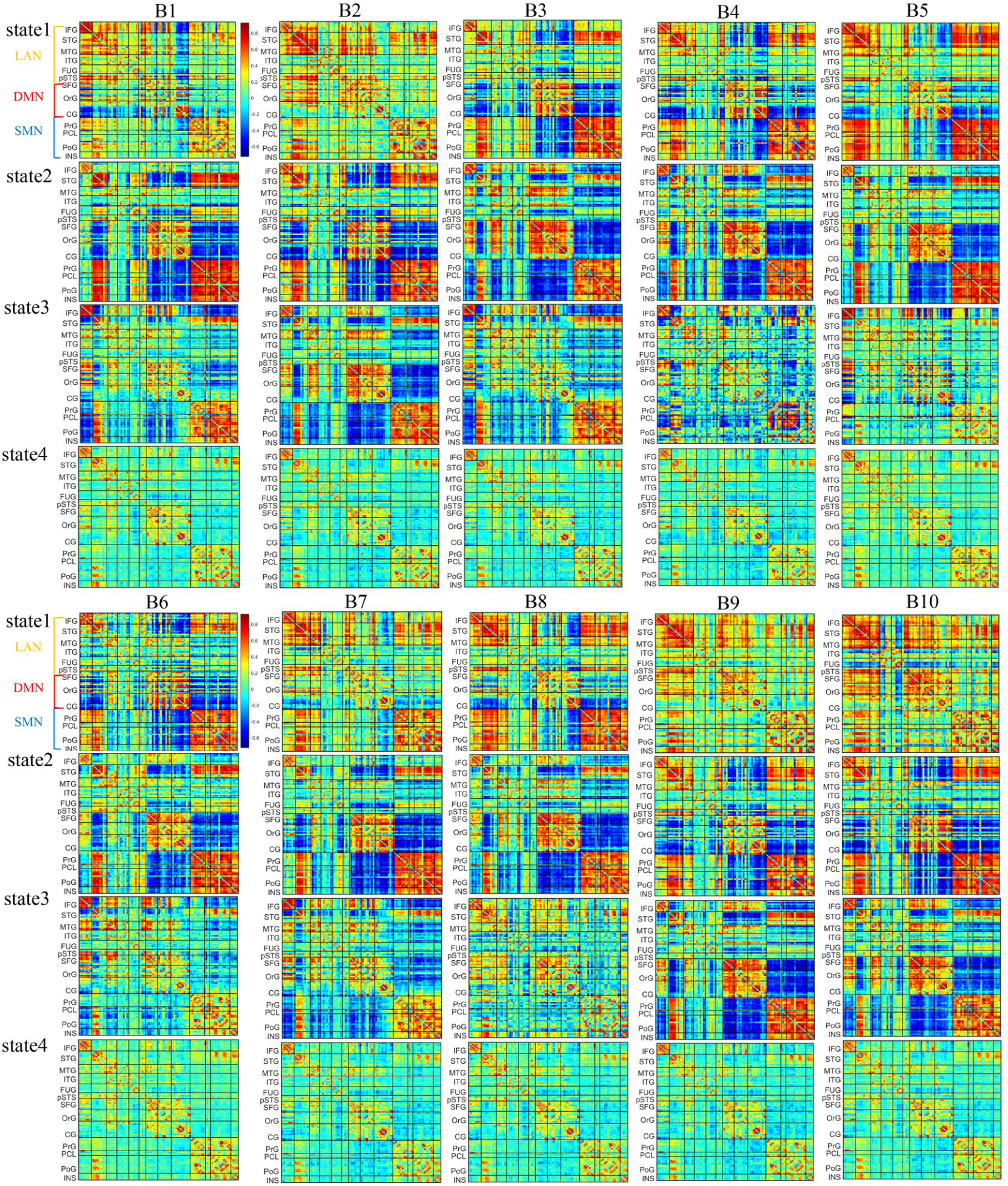
Four states identified by DCC through K-means clustering in three functional networks across background stories 1 to 10. Each functional connectivity matrix illustrates the intensity of functional connections between various nodes. (B:background)

**Figure 10.**
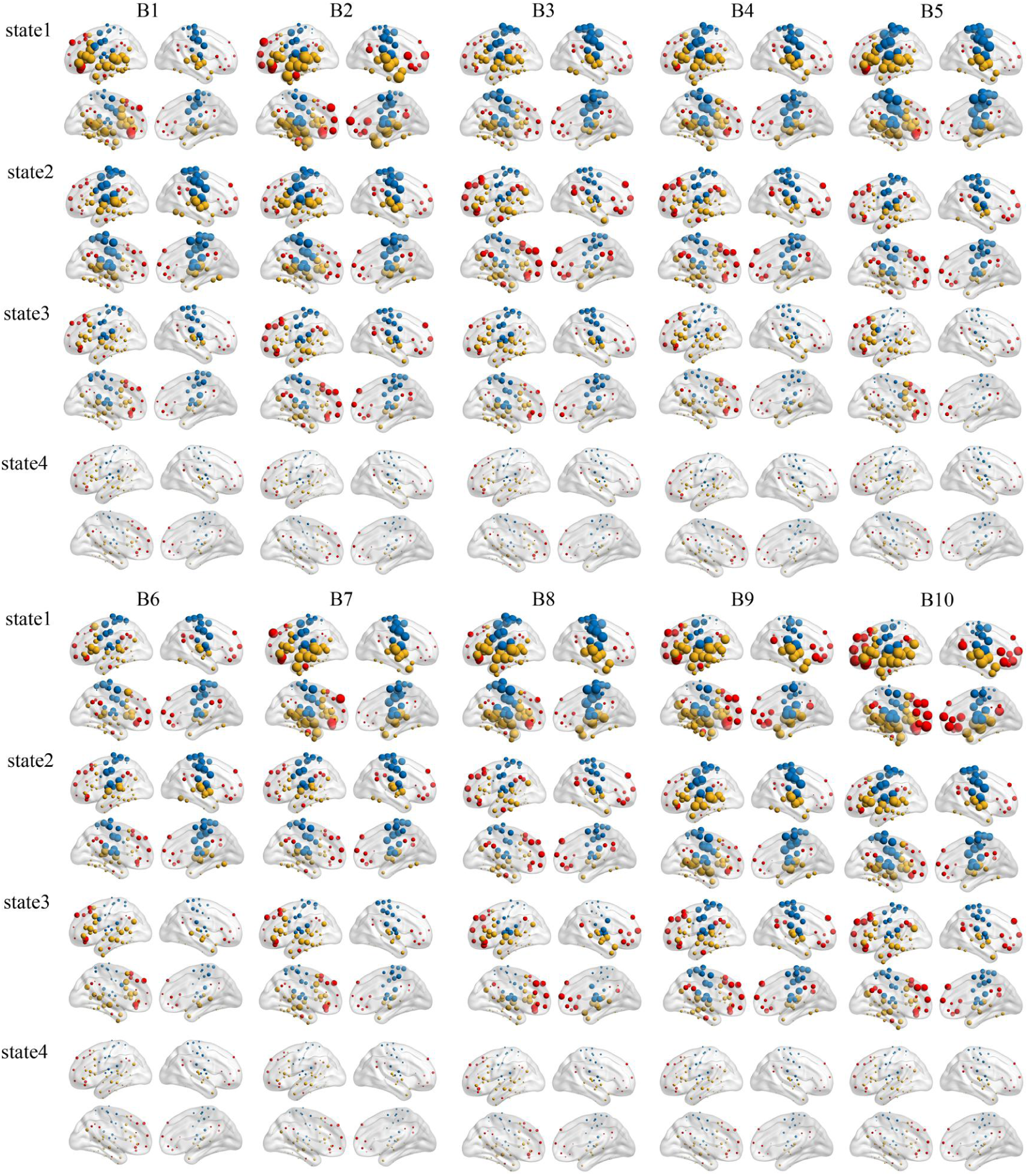
Activation node degree of DCC in three functional networks across background stories 1 to 10. The color of each node indicates its affiliation with different brain networks (blue: sensorimotor network; yellow: language network; red: default mode network), while the size of the node represents its activation intensity. (B:background)

## 4 Discussion

This study employed a naturalistic inner speech paradigm to investigate the dynamic network reorganization during situationally simulated inner speech. Participants were tasked with mentally embedding 61 cue words into specific contextual backgrounds to covertly construct stories. CAP analysis employing the ventral part of sensorimotor areas as seed points revealed predominant co-activation of the language and sensorimotor networks. CAP analysis using STG as seed points uncovered co-activation of the language, sensorimotor, and default mode networks. Dynamic conditional correlation analysis unveiled four temporal reoccurring states among these three networks. Notably, strong and persistent connections among the IFG, STG, and vPreCG were observed, indicating their sustained engagement in inner speech.

Analyses of activation during word and number inner speech, alongside term-based meta-analyses, consistently underscored the significant involvement of the bilateral vPreCG and vPoCG in inner speech. Prior studies have highlighted the crucial roles of the ventral parts of the PrG and PoG in both speech perception (Franken et al., 2022) and production (J. Lu et al., 2021). Notably, the ventral precentral gyri have been implicated as central areas for stimulation-induced speech arrest (Tate et al., 2014; Wu et al., 2015). It is theorized to serve as a “speech sound map” or “syllable table” in speech production models, potentially housing motor programs implicated in the motor planning phase of speech production (Guenther, 2016; Indefrey, 2011). Simultaneously, our findings can be interpreted through the dual-mechanism model recently proposed by, Pratts et al. This model suggests a two-dimensional cognitive framework of egocentricity and spontaneity, positing that the more egocentric and deliberate inner speech, the more it relies on articulation-derived corollary discharge (Pratts et al., 2023). In the situationally simulated inner speech tasks of the present study, participants were first presented with a background context beginning with “imagine you are…” . This approach leans towards egocentricity and deliberation, thus relying more on articulation-derived corollary discharge, which inner speech tasks have been shown to activate brain regions associated with speech production (Lurito et al., 2000; Aleman et al., 2005; Shergill et al., 2001). Our findings suggest that, apart from not involving the final articulatory execution stage, inner speech shares similarities or partial overlaps with the actual speech production process. These findings lend further support to the ConDialInt model, which posits that extended inner speech involves speech production processes up to articulatory planning, generating predictive signals with auditory characteristics, namely inner speech (Grandchamp et al., 2019).

Term-based meta-analyses also underscored the robust involvement of the middle part of the bilateral superior temporal gyrus (STG) in inner speech. While the anterior part of the STG is pivotal for auditory ventral stream functions related to word form recognition, the middle and posterior parts of the STG are crucial regions of the auditory dorsal stream involved in speech monitoring and motor control (DeWitt & Rauschecker, 2013). Furthermore, the posterior STG (pSTG) and superior temporal sulcus (STS), core regions of Wernicke’s area, are integral for lexical-phonological representations (Binder, 2017; Zhang et al., 2021). Previous research has indicated that imagining articulation activates superior temporal regions responsible for sound reconstruction (Tian et al., 2016). The ConDialInt model further posits that the STG is implicated in the formulation process, encompassing prosodic, syntactic, and morpho-phonological encoding (Grandchamp et al., 2019).

Consistent with prior investigations of inner speech (Pratts et al., 2023; Grandchamp et al., 2016; Grandchamp et al., 2019), situationally simulated inner speech elicited activation in both the language network and the sensorimotor network, indicating involvement in sensory-motor processes associated with pronunciation and sensory representations. Engaging in situationally simulated inner speech necessitates comprehension of the contextual background story and the word information presented. This entails word recognition, grammatical structure analysis, sentence interpretation, and semantic comprehension. Furthermore, participants must select appropriate vocabulary and construct complex sentences for mental narration, even though this process may be condensed at the syntactic, lexical, and phonological levels (Fernyhough, 2004; Geva et al., 2011; Grandchamp et al., 2019; Vygotsky, 2012).

Numerous studies have demonstrated the involvement of the sensorimotor network (SMN) in memory retrieval transformation and imagery construction (Kleber et al., 2007; Lima et al., 2015; Tian et al., 2016). It plays a pivotal role in general sequence processing across various domains (Cona & Semenza, 2017) and is activated during timing and initiation in speech motor control, as well as top-down prediction during speech perception and construction (Hertrich et al., 2016; Lu et al., 2023). Participants must vividly construct scenarios and scenes in their minds and create verbal expressions to convey the narrative content. The SMN is crucial for processes related to sensory simulation, motor control, and planning (Ionta et al., 2011; Tian & Poeppel, 2010; Wolpert & Flanagan, 2001). Its activation may facilitate internal generation of sensory experiences, motor planning, and scenario simulations relevant to situationally simulated inner speech.

Our results demonstrate for the first time that the default network (DMN) is involved in processing inner speech. This provides evidence for the involvement of inner speech in aspects such as memory, self-observation, introspection, and metacognition. Traditionally, the DMN has been regarded as an “intrinsic” system that is suppressed during tasks demanding external attention (Buckner et al., 1995;

Buckner, 2012; Buckner & DiNicola, 2019; Raichle et al., 2001; Shulman et al., 1997). However, recent evidence suggests that the DMN serves as an active and dynamic ‘sense-making’ network, facilitating the integration of extrinsic and intrinsic information over long timescales (Yeshurun et al., 2021a). In situationally simulated inner speech tasks, participants are tasked with internally integrating background stories with upcoming word events, relying on long-term memory, patterns, and belief systems to internally generate vivid speech. The involvement of the DMN in situationally simulated inner speech may mediate the retrieval and integration of such personalized representations. Given that the story backgrounds and word presentations are randomized, the variability in DMN responses among participants may be attributed to personalized prior biases and differences in story content. The absence of significant DMN activation in the co-activation pattern of the bilateral vPreCG and vPoCG may be due to their roles in the final generation stage of inner speech.

Furthermore, in seed-based co-activation pattern analysis, extensive activation of cerebellar regions was observed. Numerous studies have provided compelling evidence that the cerebellum plays a crucial role in cognitive and affective functions beyond its historical association with motor function (Diedrichsen et al., 2019; Schmahmann et al., 2019; Strick et al., 2009). Specifically, at the Crus I/II apex, regions associated with cerebral networks involved in functions such as language, memory, and social cognition have been identified (Gaiser et al., 2024). Our results indicate that the process of situationally simulated inner speech activates the language network located at the edge of the cerebellar lobules I/II and the default network within the cerebellum (King et al., 2019; Xue et al., 2021). According to the multi-domain task battery parcellation (Gaiser et al., 2024), regions related to the default network can be subdivided into different areas related to narrative comprehension (regions 7 and 8), language functions (regions 8 and 9), and autobiographical recall (region 10) (King et al., 2019). The activation of the right cerebellum in our results aligns with the ConDialInt model, where speech targets are sent to the cerebellum (Grandchamp et al., 2019). However, the current functional magnetic resonance imaging (fMRI) data are insufficient to fully assess the roles played by different parts of the cerebellum in situationally simulated inner speech.

For network reorganization in situationally simulated inner speech, it is evident that interactions among the three networks change over time. Moreover, we consistently observed strong connections among the inferior frontal gyrus (IFG), superior temporal gyrus (STG), and precentral gyrus (PrCG) across all four states, indicating consistent activity in these brain regions, which aligns with findings from meta-analyses and CAP results.

To examine whether contextual backgrounds could be decoded from network reorganizations, we employed the support vector machine (SVM) algorithm for multiclass classification analysis. However, average decoding accuracies of multi-label SVM based on two networks (SMN and LAN) and three networks (SMN, LAN and DMN) are 11% and 10% respectively, which closely resemble chance level (Supplementary figure 16). We attempt to provide some explanations for this. Our findings indicate that the primary regions associated with inner speech are situated in the speech perception and motor areas. While the activation of these regions is essential for the functionality of speech perception and articulatory systems during inner speech, their contribution to semantic processing may be limited. Consequently, these regions may have a restricted role in decoding the semantic information of inner speech. Another potential factor could be the insufficient discriminability of experimental stimuli in terms of semantics, lacking robust and systematic semantic distinctions. This observation is consistent with the findings indicating poor inter-subject consistency in CAP and DCC.

This study is a preliminary exploration and has some limitations. Firstly, due to limitations in publicly available datasets and the lack of clear behavioral manifestations in situationally simulated inner speech, it is challenging to explain the behavioral significance of network reorganization. Subsequent studies could optimize experimental design by having participants recall and subjectively rate the vividness and continuity of their imagined content after the experiment (Hurlburt et al., 2013; Lima et al., 2015). Secondly, while the experimental design using randomized stimuli controlled certain variables, it also introduced inconsistency in participants’ responses, making it challenging to analyze the data using inter-subject analysis (Hasson et al., 2004; Lerner et al., 2011; Nastase et al., 2019; Nastase et al., 2021), and rendering SVM decoding ineffective. Future research could improve the experimental design to better understand the behavioral implications of network reorganization. Additionally, the stimuli used in the study were limited to abstract words and did not include concrete words. There are differences in the representation of concrete and abstract words, with a greater involvement of the language system in processing abstract concepts, and a greater involvement of the perceptual system in processing concrete concepts (Wang et al., 2010). Future research could investigate the impact of abstractness on situationally simulated inner speech. Overall, this study provides initial insights into situationally simulated inner speech, but further research is needed to gain a deeper understanding of its nature.

## 5 Conclusions

Situationally simulated inner speech, as a unique yet common manifestation of inner speech, finds its core brain area in the bilateral ventral portion of the sensorimotor cortex. Additionally, the middle portion of the superior temporal gyri may serve as another core brain region. As a dynamic and multi-cognitive processing task, situationally simulated inner speech necessitates the involvement of the language network for a truncated form of overt speech, sensorimotor network for perceptual simulation and monitoring, and default mode network for integration and ‘sense-making’ processing.

