## Supplementary materials for "How do we imagine a speech? A triple network model for situationally simulated inner speech"

**How do we imagine a speech? Dynamic network reorganization during situationally simulated inner speech**

**Supplementary materials**

**Supplementary Tables**

**Supplementary Table 1. English translations for the 61 cue words used in the experiment.**

| development | profit | law | home |
| --- | --- | --- | --- |
| politics | end | possibility | damage |
| beginning | movement | punishment | competition |
| goal | growth | interest | quality |
| help | peace | fight | reform |
| decision | error | collaboration | truth |
| cost | critique | anxiety | foundation |
| success | value | safety | agreement |
| future | death | might | verdict |
| problem | defiance | task | culture |
| information | opinion | discussion | border |
| doubt | love | danger | age |
| fortune | victory | argument | hope |
| demand | closeness | past | opposition |
| technique | leadership | freedom | sense |
| force |  |  |  |

**Supplementary Table 2. English translations for the 10 background contexts used in the experiment.**

| Imagine you are an astronaut exploring faraway galaxies. There you meet members of a previously unknown alien race. Try to explain humanity and its culture to them. | Imagine you are the best man / maid of honor at your best friend’s wedding. At the party, you are giving a speech, in which you recall the past and tell a few anecdotes from your common past. |
| --- | --- |
| Imagine you are an adventurer on your way to the north pole. You are writing a chapter of your travel diary, in which you emphasize your willingness to successfully bring the expedition to an end, despite low supplies and the bitter cold. | Imagine your partner finds text messages with a potential lover on your mobile phone. You get into an argument and you are trying to explain yourself. |
| Imagine you are a member of the parliament from the Green party. You are  giving a speech about the challenges posed by climate change and the right measures to stop a catastrophic global warming. | Imagine you are a defense attorney, and you are representing a bank robber at court. You are making a summation for the accused, who committed the robbery to be able to pay for a life-saving surgery for his wife. |
| Imagine you are part of a group of cave explorers. You will be the first humans to explore a newly discovered stone-age cave. Prior to the expedition, you speak to a tv team and explain to them what the potential discovery of new cave paintings would mean for understanding early human history. | Imagine you are an unrenowned runner who totally surprisingly finishes the Berlin marathon in second place. Afterwards, the media accuse you of doping. In a press statement, you try to convince the public that you are innocent. |
| Imagine you are a German diplomat on your way to an important EU summit. There, you will be asked to outline political measures you consider suitable for containing the “refugee crisis”. | Imagine you are a social democrat politician in 19th century Germany. You get the chance to speak to the Kaiser and convince him to strengthen the rights of the parliament to create a more modern and fairer political system. |

**Supplementary Figures**

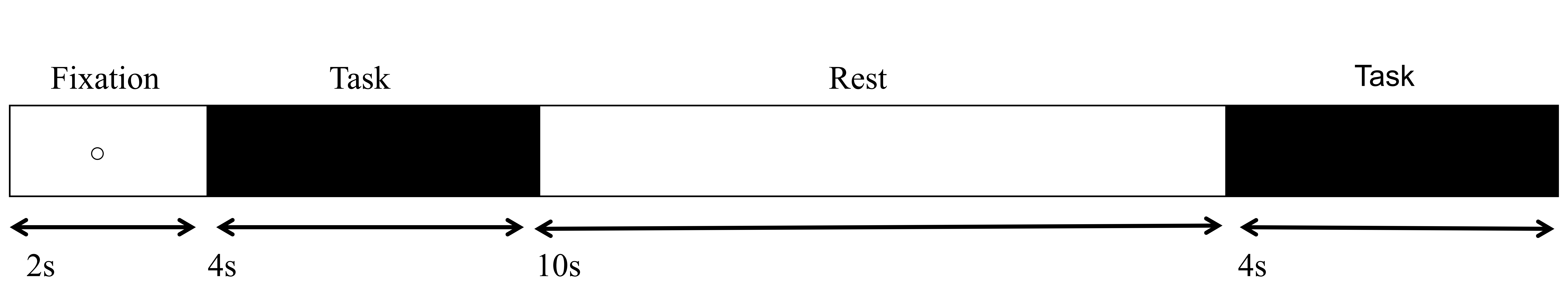

Supplementary figure 1. fMRI experimental procedure in the "Bimodal" Dataset

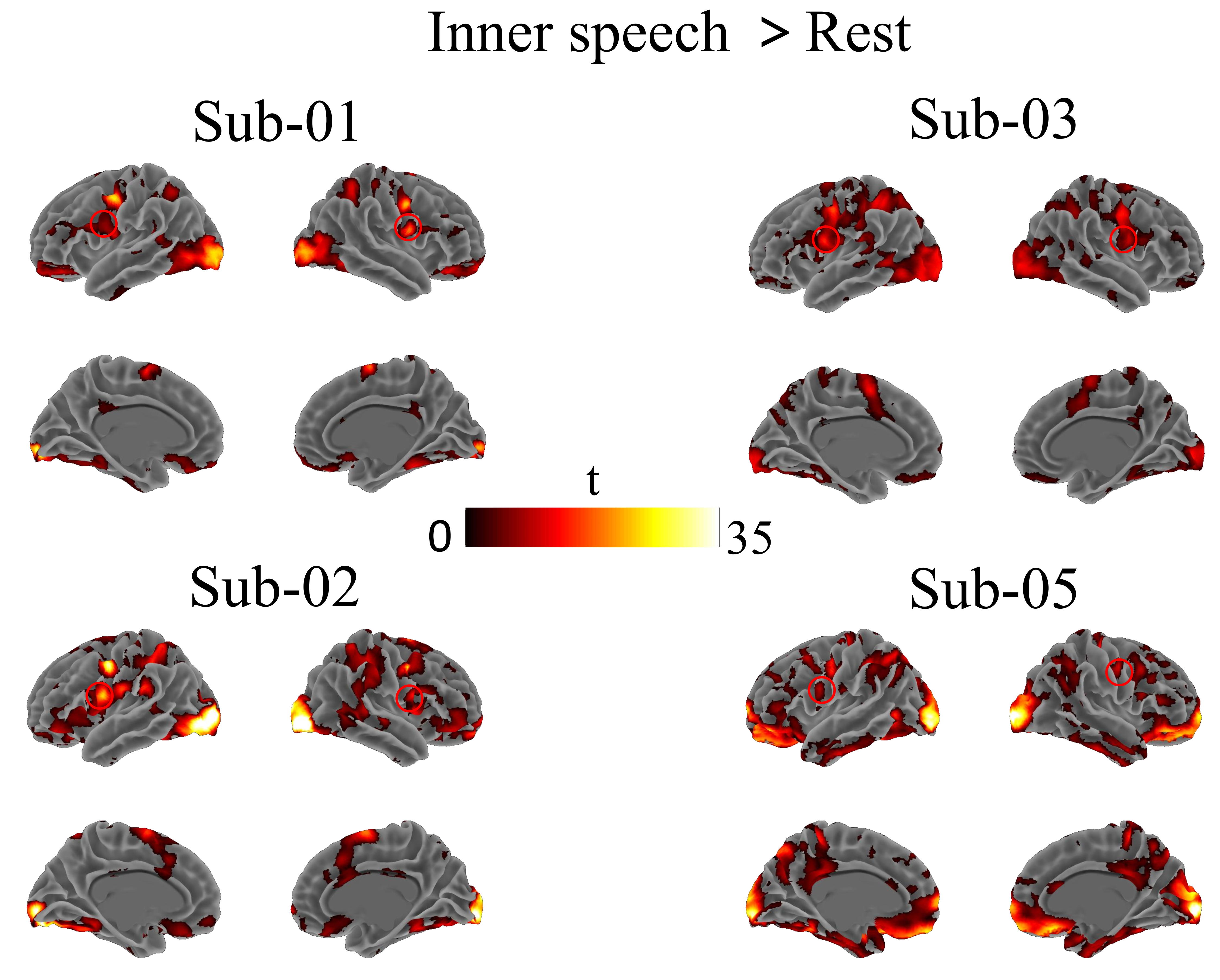

Supplementary figure 2. fMRI activations for covert speech of words and numbers. Voxel-wise thresholding was conducted with FDR correction, using a voxel-level threshold of cluster size > 20, and corrected p < 0.05.

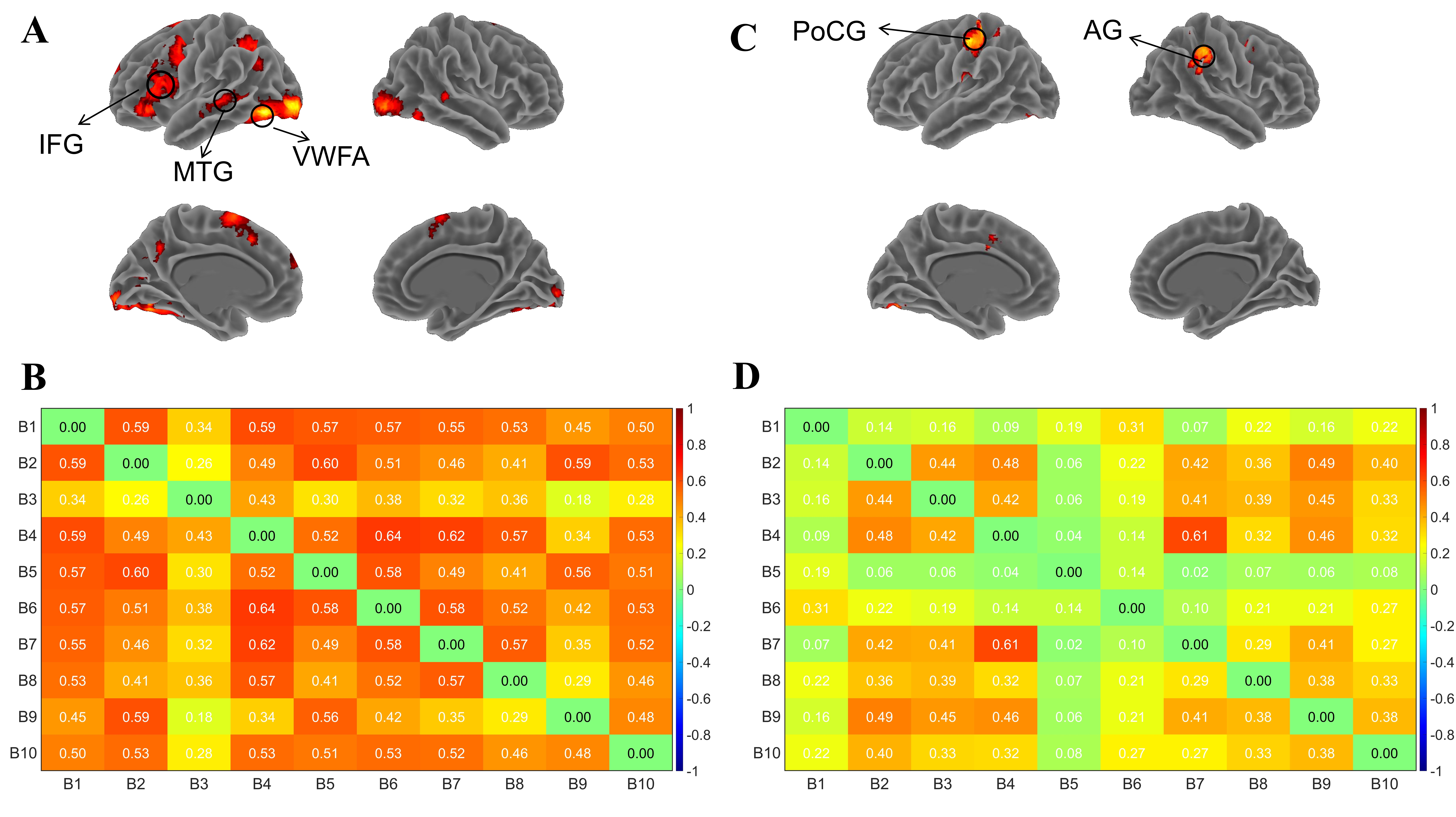

Supplementary figure 3. Brain activations of visual word processing (A) and left finger button press (C) and their spatial similarities across the 10 runs (B & D). FDR correction at p < 0.05 with cluster size of 20. To prevent samples from being matched with themselves and to simplify the matrix representation, we set the diagonal values of the similarity matrix to 0.

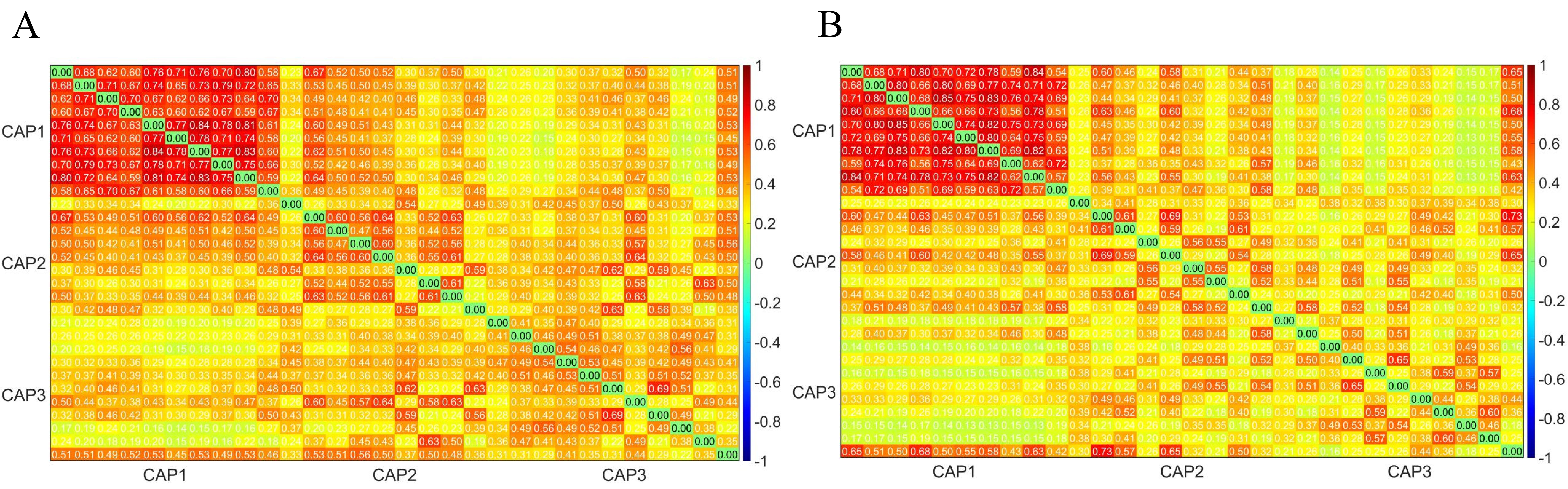

Supplementary figure 4. Spatial similarity of three CAPs with left vPreCG+vPoCG (A) and the right vPreCG+vPoCG (B) as the seeds across 10 runs. Modules within the same network (along the diagonal) contain the similarity between the 10 background stories. To prevent samples from being matched with themselves and to simplify the matrix representation, we set the diagonal values of the similarity matrix to 0.

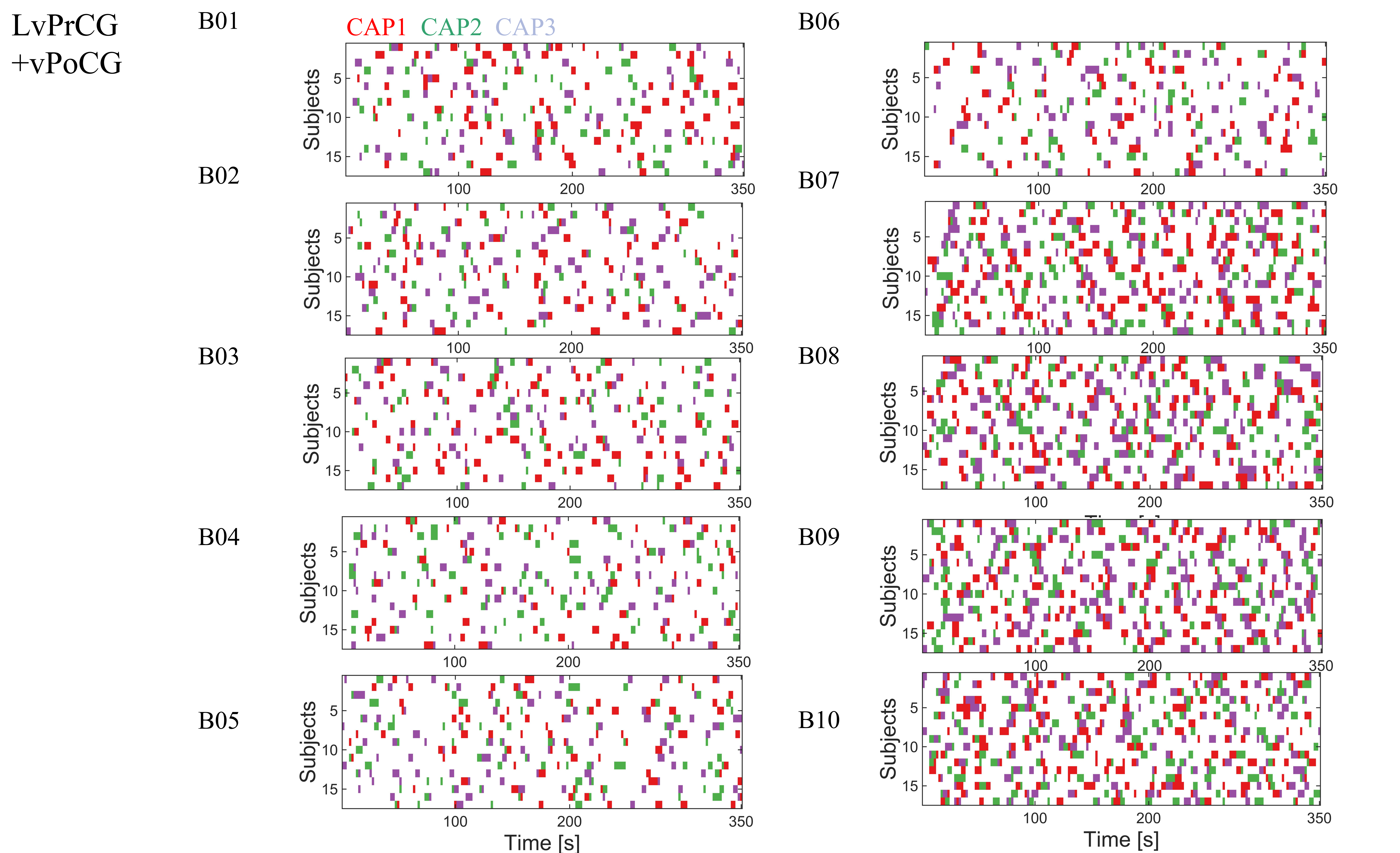

Supplementary figure 5. The transition vectors of each participant over time in the situationally simulated inner speech task across the 10 background stories, using the ventral part of the left precentral gyrus and postcentral gyrus as seed points.

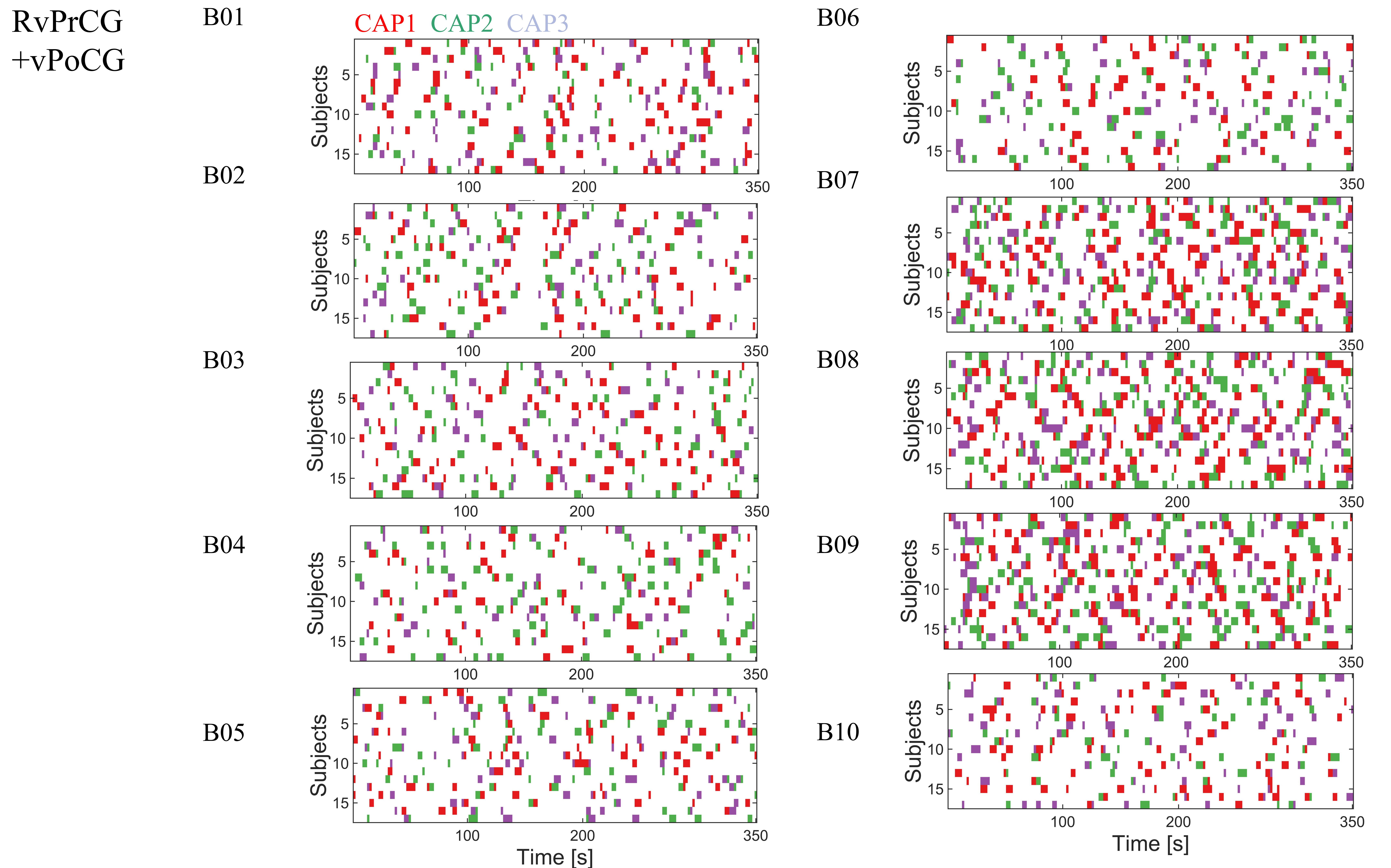

Supplementary figure 6. The transition vectors of each participant over time in the situationally simulated inner speech task across the 10 background stories, using the ventral part of the right precentral gyrus and postcentral gyrus as seed points.

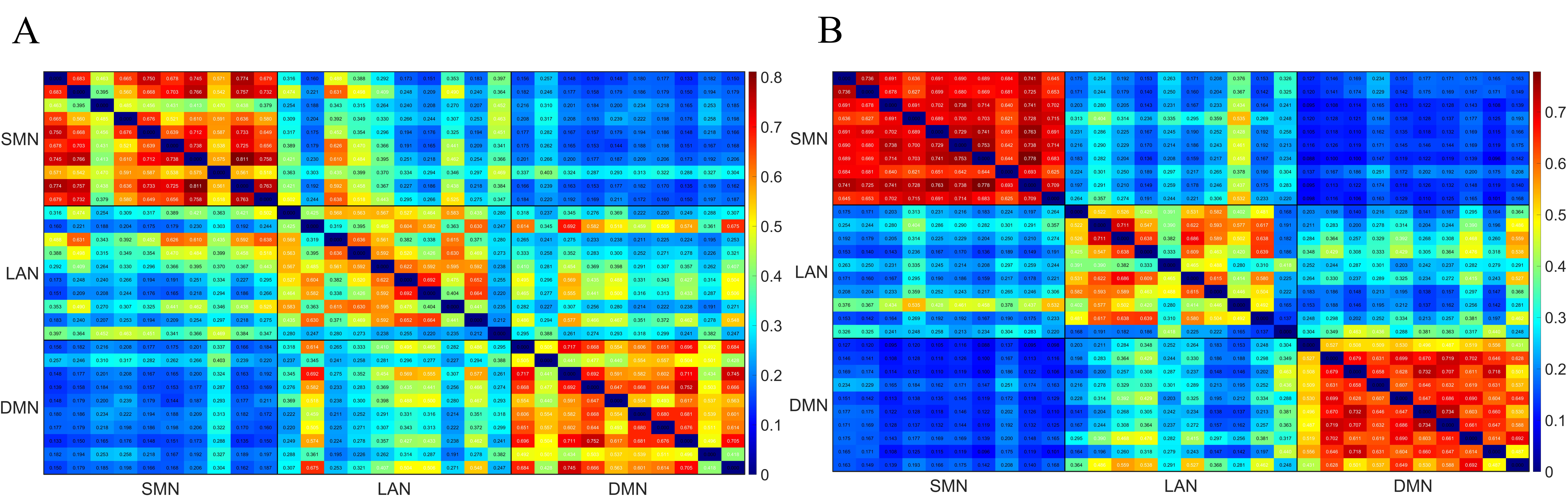

Supplementary figure 7. Spatial similarity of three CAPs with the LSTG (A) and RSTG (B)as the seedS across 10 runs. Modules within the same network (along the diagonal) contain the similarity between the 10 background stories. To prevent samples from being matched with themselves and to simplify the matrix representation, we set the diagonal values of the similarity matrix to 0.

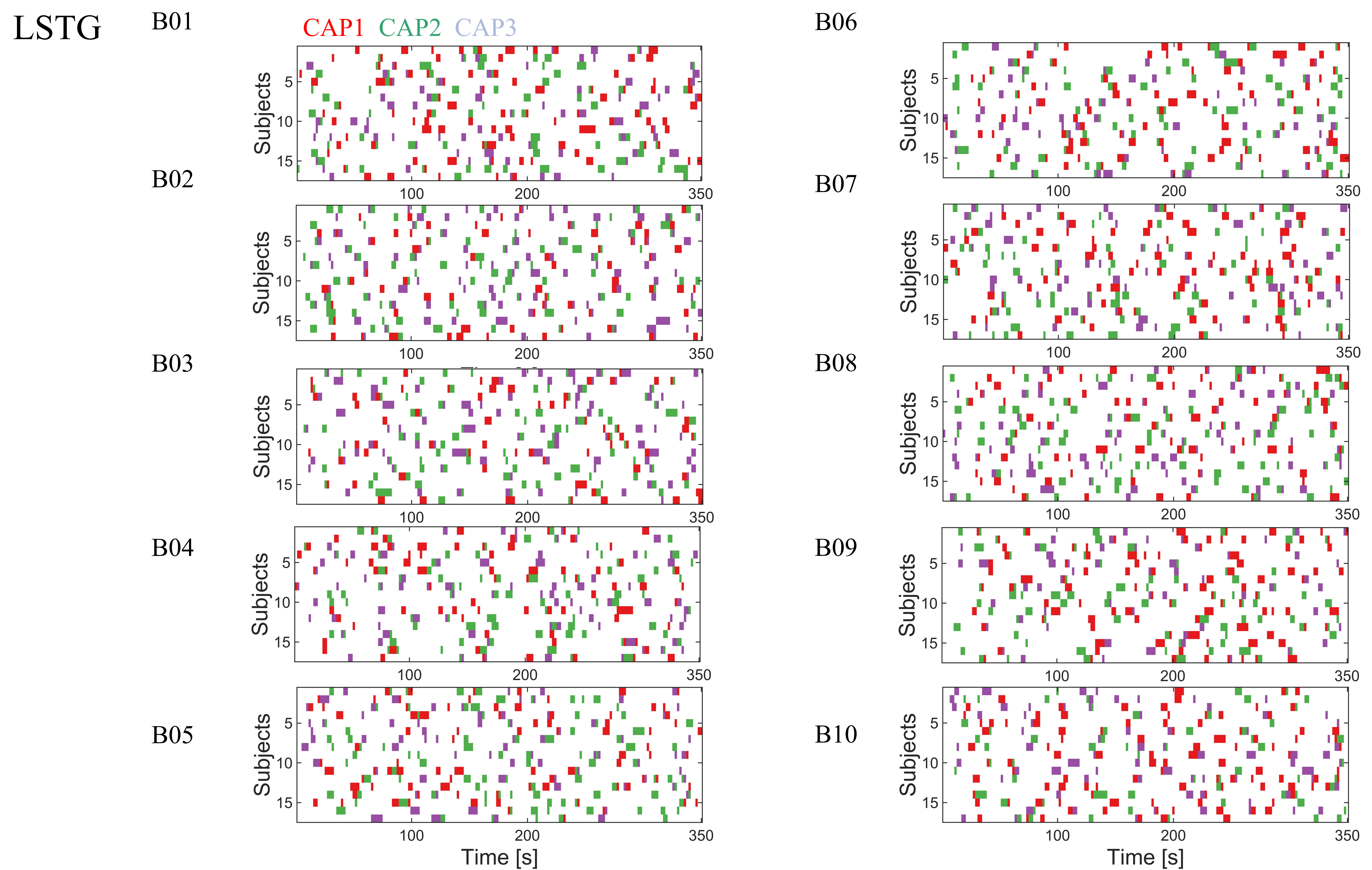

Supplementary figure 8. Transition vectors of each participant over time in the situationally simulated inner speech task across the 10 background stories, using the left superior temporal gyrus as the seed point.

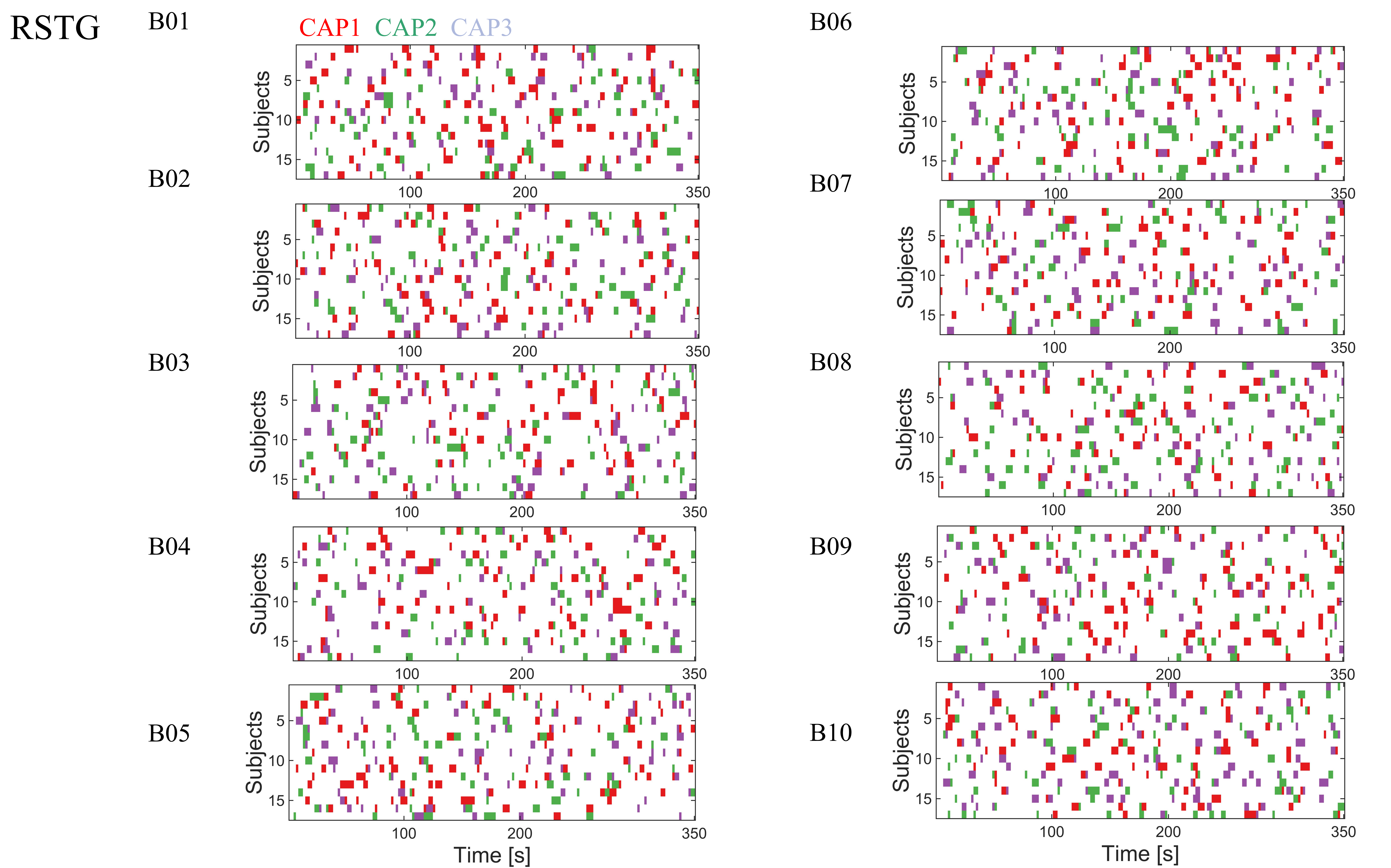

Supplementary figure 9. Transition vectors of each participant over time in the situationally simulated inner speech task across the 10 background stories, using the right superior temporal gyrus as the seed point.

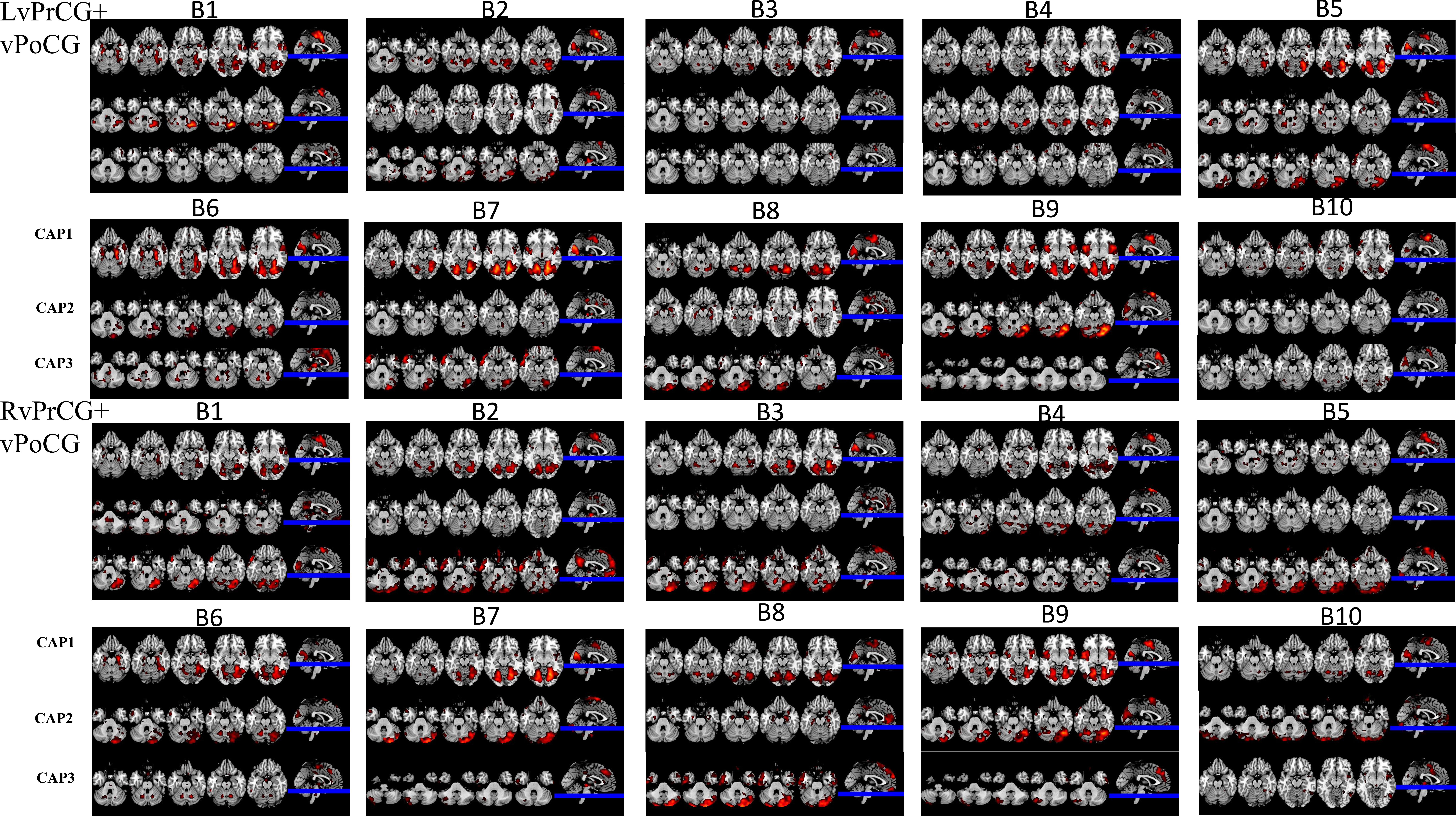

Supplementary figure 10. Activation of the cerebellum in the ventral parts of the bilateral precentral and postcentral gyrus across Background Stories 1 to 10 (Z > 1, cluster size = 20 voxels).

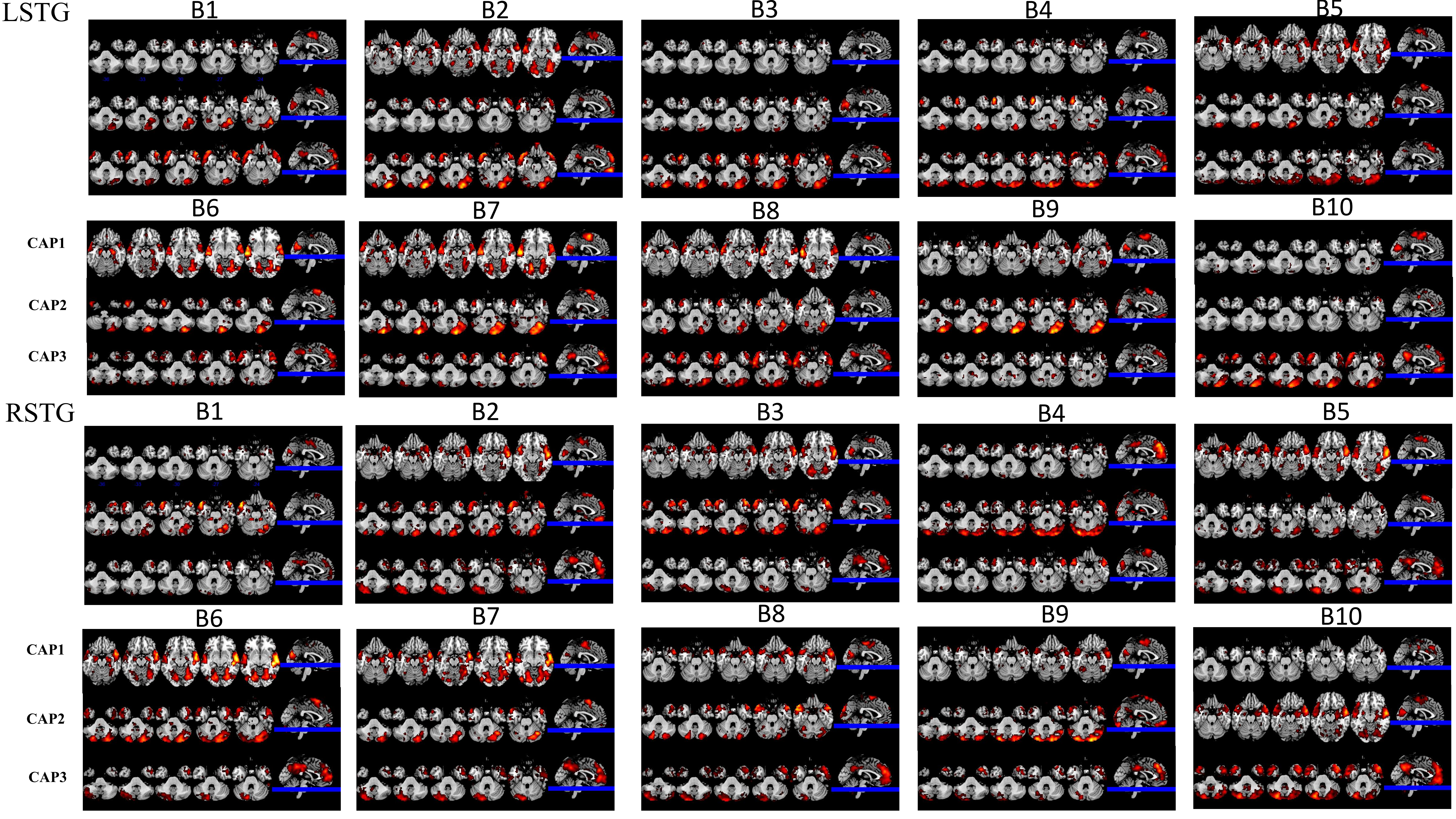

Supplementary figure 11. Activation of the cerebellum by bilateral superior temporal gyrus across Background Stories 1 to 10 (Z > 1, cluster size = 20 voxels).

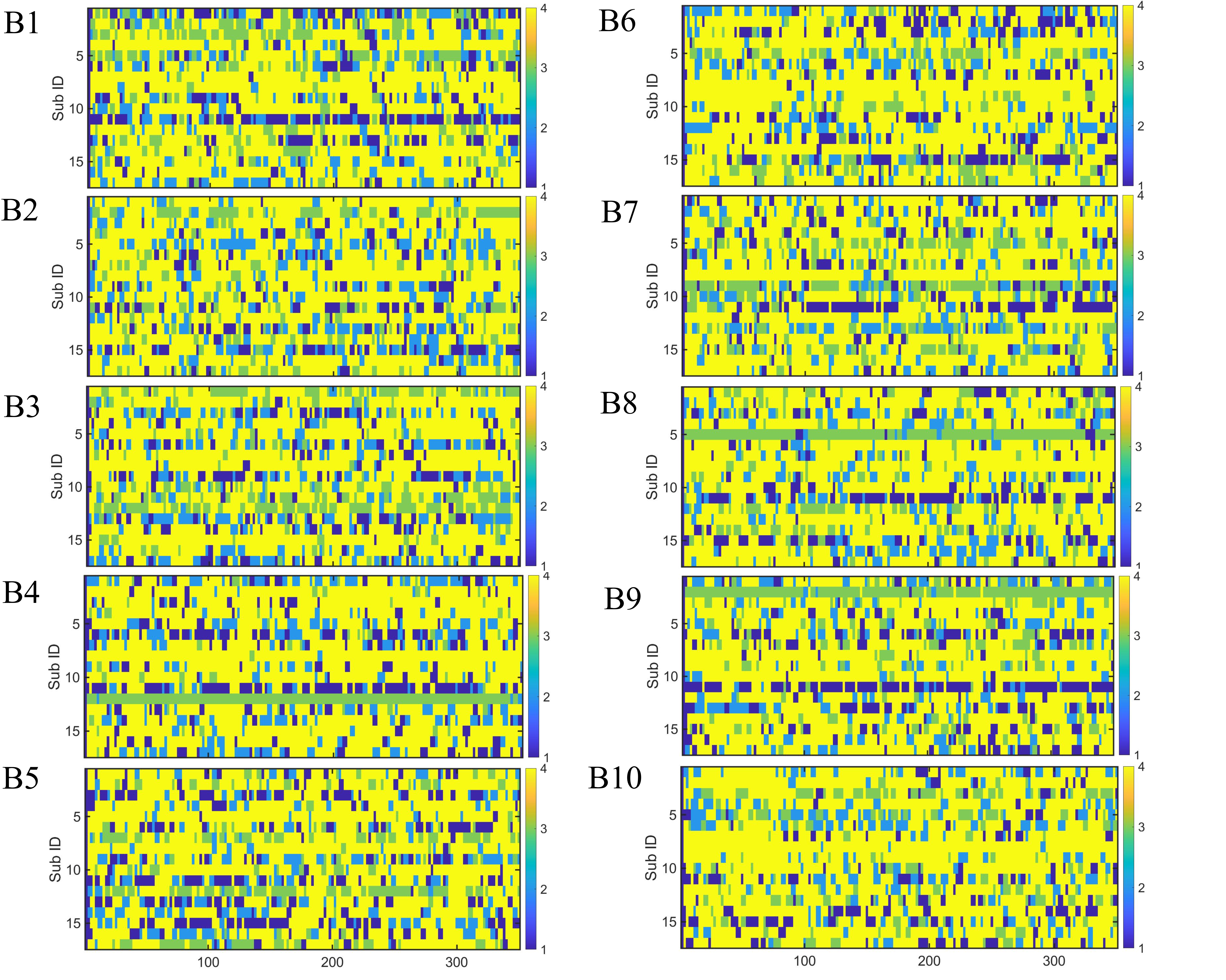

Supplementary figure 12. State transition matrix of DCC activation in two functional networks across background stories 1 to 10.

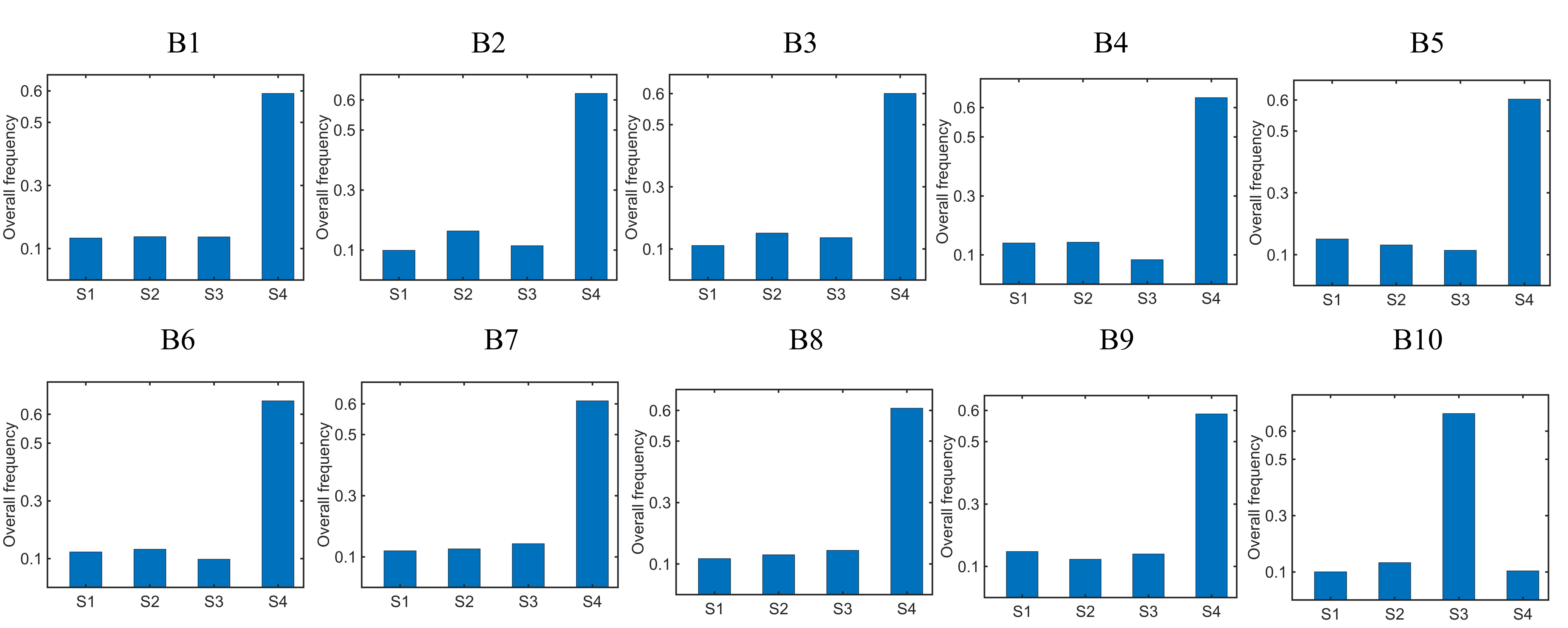

Supplementary figure 13. Overall frequency of the four states activated by DCC in two functional networks across background stories 1 to 10.

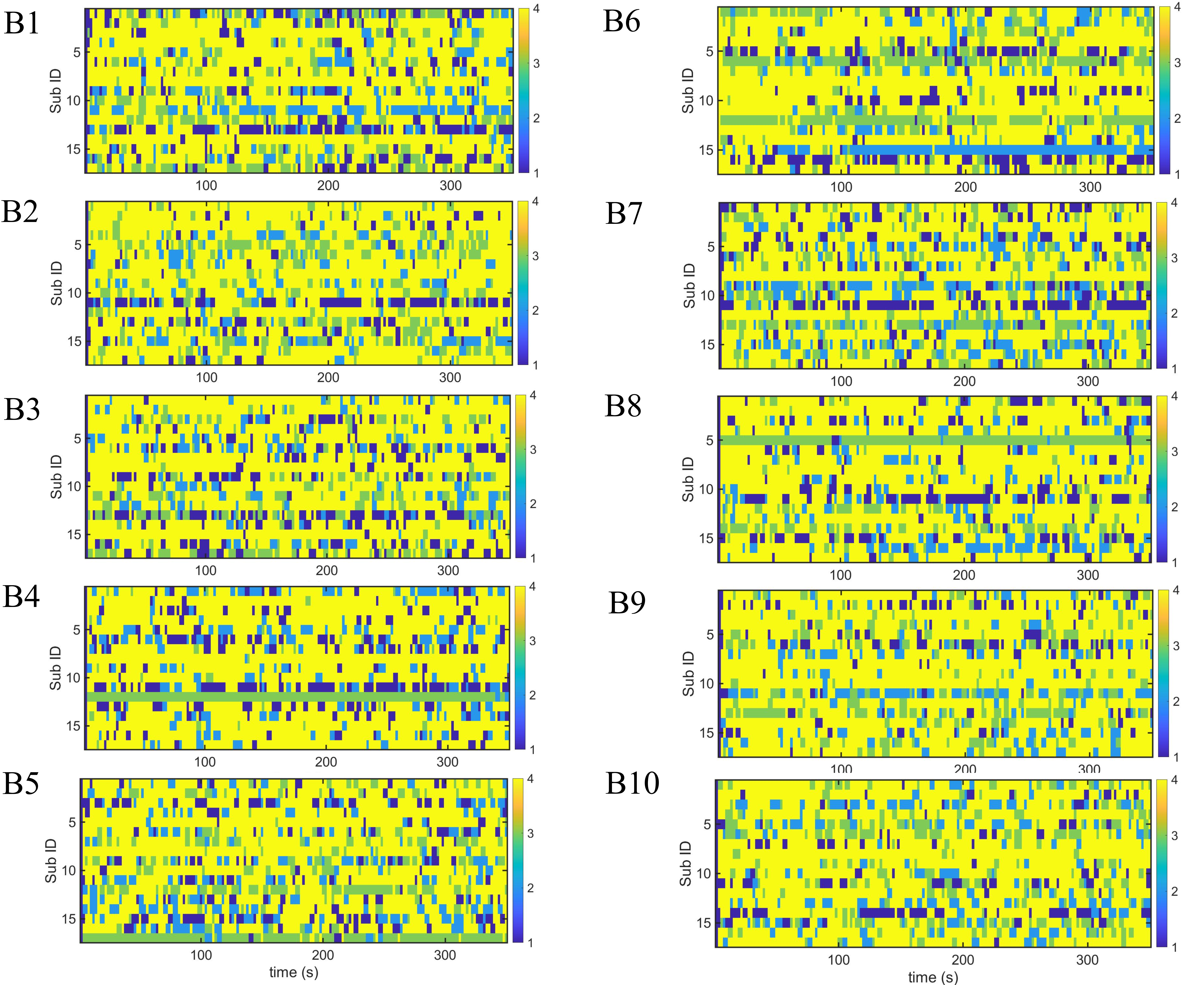

Supplementary figure 14. State transition matrix of DCC activation in three functional networks across background stories 1 to 10.

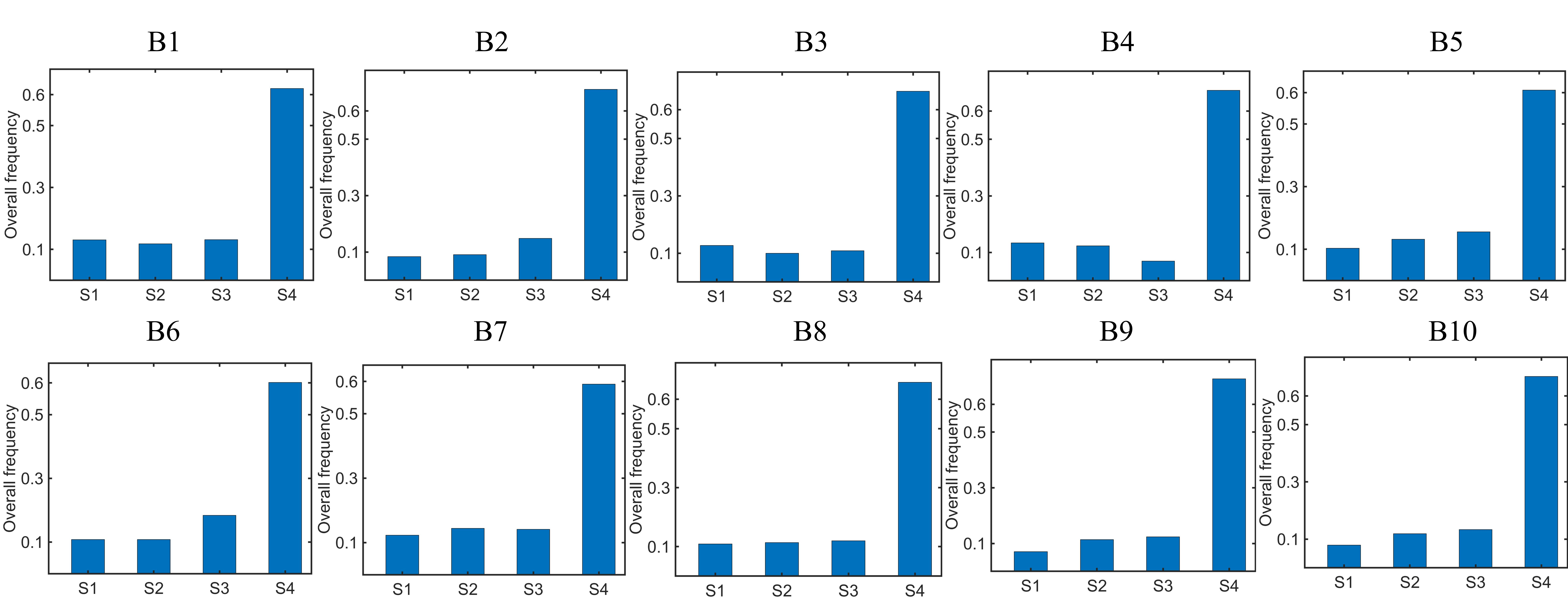

Supplementary figure 15. Overall frequency of the four states activated by DCC in three functional networks across background stories 1 to 10.

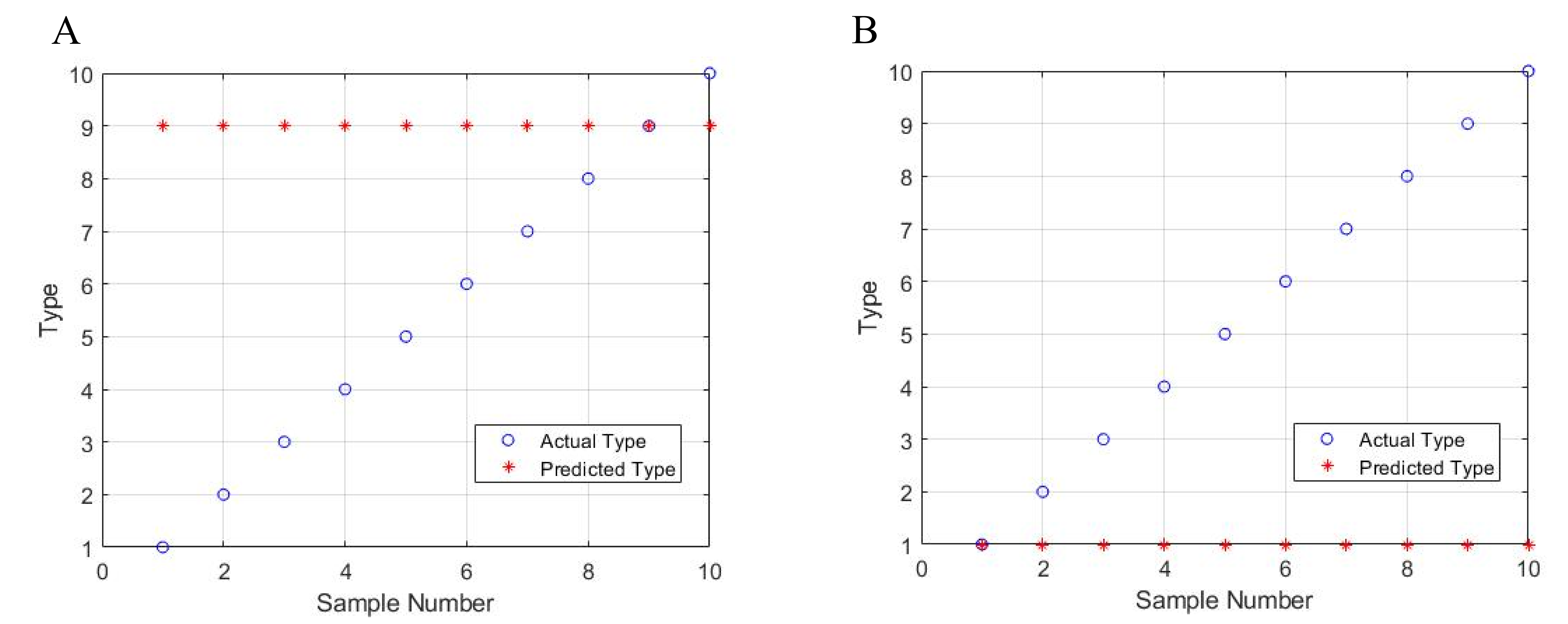

Supplementary figure 16. The average decoding accuracies of multi-label SVM based on two networks (SMN and LAN) and three networks (SMN, LAN and DMN) are 11.76% (A) and 10% (B) respectively.
